# Caterpillars on a phytochemical landscape: the case of alfalfa and the Melissa blue butterfly

**DOI:** 10.1101/498139

**Authors:** Matthew L. Forister, Su’ad Yoon, Casey S. Philbin, Craig D. Dodson, Bret Hart, Joshua G. Harrison, Oren Shelef, James A. Fordyce, Zachary H. Marion, Christopher C. Nice, Lora A. Richards, C. Alex Buerkle, Zach Gompert

## Abstract

Modern metabolomic approaches that generate more comprehensive phytochemical profiles than were previously available are providing new opportunities for understanding plant-animal interactions. Specifically, we can characterize the phytochemical landscape by asking how a larger number of individual compounds affect herbivores and how compounds covary among plants. Here we use the recent colonization of alfalfa (*Medicago sativa*) by the Melissa blue butterfly (*Lycaeides melissa*) to quantify plant metabolites and the performance of caterpillars as affected by both individual compounds and suites of covarying phytochemicals. We find that survival, development time and adult weight are all associated with variation in nutrition and toxicity, including biomolecules associated with plant cell function as well as putative anti-herbivore action. The plant-insect interface is complex, with clusters of covarying compounds in many cases encompassing divergent effects on different aspects of caterpillar performance. Individual compounds with the strongest associations are largely specialized metabolites, including alkaloids, phenolic glycosides and saponins. The saponins are represented in our data by more than 25 individual compounds with beneficial and detrimental effects on *L. melissa* caterpillars, which highlights the value of metabolomic data as opposed to approaches that rely on total concentrations within broad defensive classes.

## 1 INTRODUCTION

One of the conceptual pillars of trophic ecology is the idea that herbivores must overcome the barrier of plant defensive chemistry before extracting the nutrients necessary for growth and reproduction (Feeny, Rosenthal, & Berenbaum, 1992). The success of this idea is reflected in areas of research that include coevolution (Agrawal, Petschenka, Bingham, Weber, & Rasmann, 2012), ecological specialization (Dyer, 1995), and nutrient flow in ecosystems (Hättenschwiler & Vitousek, 2000). In most cases, progress has been made by chemical ecologists focusing on small subsets of the specialized metabolites produced by plants and consumed by herbivores. The focus on a few charismatic molecules or classes of compounds, such as furanocoumarins (Berenbaum, 1983) or cardiac glycosides (Zalucki, Brower, & Alonso-M, 2001), was at least in part necessitated by early methods in natural products chemistry that were targeted and not easily optimized for the discovery of large suites of co-occurring metabolites (Dyer et al., 2018; Maag, Erb, & Glauser, 2015). As technological limitations have dissipated, the opportunity now exists for a more comprehensive understanding of the challenges faced by herbivores, with the possibility of discovering, among other things, novel compounds and synergistic interactions among compounds (Prince & Pohnert, 2010; Richards, Dyer, Smilanich, & Dodson, 2010; Sardans, Penuelas, & Rivas-Ubach, 2011). More generally, an important task is to quantify the phytochemical complexity of the antagonistic interaction between plants and herbivores, with an eye towards understanding constraints on the evolution of both players (Fordyce & Nice, 2008; Macel, van Dam, & Keurentjes, 2010) and predicting the formation of new plant-herbivore interactions (Erbilgin, 2018). Here we use the example of a specialized herbivore and a recently colonized host plant to investigate the phytochemical landscape from the perspective of developing caterpillars. By the “phytochemical landscape” we mean metabolomic variation among individual plants and associated toxic and nutritional effects on, in our case, a focal herbivore (Glassmire et al., 2019; Hunter, 2016; Wu, Wilson, Chang, & Tian, 2019).

The Melissa blue butterfly, *Lycaeides melissa*, is specialized on larval host plants in the pea family (Fabaceae), primarily in the genera *Astragalus* and *Lupinus*. Within the last 200 years, *L. melissa* has colonized introduced alfalfa, *Medicago sativa* (Fabaceae), at least twice and probably multiple times (Chaturvedi et al., 2018), forming a heterogeneous patchwork of association throughout the range of the butterfly in western North America, often with naturalized or weedy patches of *M. sativa*. In general, *M. sativa* is a suboptimal host plant for *L. melissa*: individuals that feed on the plant have reduced survival and performance relative to individuals feeding on native hosts (Forister, Nice, Fordyce, & Gompert, 2009). *M. sativa*-associated populations do, however, show evidence for a slight increase in the ability to develop on the novel resource relative to populations that remain on native plants (Gompert et al., 2015). Additional evolutionary change in populations associated with the novel host is evidenced by reduced female oviposition preference for native hosts (Forister et al., 2012) and reduced caterpillar performance on native hosts (relative to populations that have not shifted to the exotic) (Gompert et al., 2015).

The genetic architecture of host use in this system is known to be polygenic and characterized by loci with conditionally neutral (host-specific) effects and ongoing local adaptation (Gompert et al., 2015). What is needed next is an understanding of which plant traits most affect *L. melissa* fitness. Previous work has suggested that phytochemical variation among host populations is biologically significant for caterpillars eating *M. sativa* (Harrison et al., 2016), but the magnitude of these effects and the salient compounds are unclear. Moreover, caterpillars do not encounter compounds in isolation, but in combinations of covarying molecules, and it is unknown how variation among hosts in phytochemical mixtures affects herbivore evolution. For example, will the trajectory of further local adaptation by *L. melissa* to *M. sativa* be a matter of evolving the ability to detoxify one or a large number of compounds? A better understanding of how key compounds covary among individual plants could also shed light on the potential for evolutionary response of the plant to herbivores in its introduced North American range. Here we use a common garden approach and caterpillars individually reared in a controlled environment to address these questions while describing the effects of metabolomic variation in *M. sativa* on *L. melissa*.

## 2 METHODS

### 2.1 Plants and caterpillars

Plants used in this project were grown at the University of Nevada, Reno, Main Station experimental farm. The common garden was planted in 2016 with seeds collected the previous year from 45 plants (previously studied by [Harrison et al., 2018]) growing in a fallow field in north-western Nevada on the western edge of the Great Basin Desert. The focal butterfly, *L. melissa*, was present in the source field but has not colonized the university farm where experimental plants were grown. The 45 maternal plants each contributed 15 offspring to a randomized grid design in the common garden, irrigated with broadcast sprayers in 2016 and drip in 2017, without supplemental fertilization. A single plant was randomly selected from each maternal family for use in the rearing experiment reported here as a way to capture as much genetic and phenotypic variation as possible.

On 17 and 18 July 2017, a total of 45 *L. melissa* females were collected from an alfalfa-associated population near Verdi, NV, and confined to oviposition arenas (three females per arena, 500 mL plastic cups) with host plant leaves and mesh lids sprayed with Gatorade^®^, a sports drink with sugar, water, carbohydrates, salt and other ingredients that has been used elsewhere for captive butterflies (Mattila & Otis, 2003). After three days, eggs were removed from leaves, pooled, and kept at room temperature until hatching, at which time caterpillars were placed individually in Petri dishes (100 x 25 mm) with leaves of a particular *M. sativa* individual (which became the only plant from which they were fed throughout the experiment). Ten caterpillars were assigned to each of the 45 experimental *M. sativa* plants (for a total of 450 independently-reared caterpillars) and kept in a growth chamber set to 25°C and a 12 hour light / 12 hour dark cycle. Caterpillars were given new, undamaged leaves as needed, approximately every 2-3 days. From each caterpillar we recorded survival to adult, sex, date of eclosion (if successful) and adult weight to the nearest 0.01 mg on a Mettler Toledo XP26 microbalance. Adult weight is taken as a proxy for fitness in *L. melissa* (Forister et al., 2009).

### 2.2 Phytochemistry and plant traits

Metabolomic variation among individual plants was characterized with liquid chromatography– mass spectrometry (LC-MS) (Jorge, Mata, & António, 2016) using leaves collected on a single day at the start of the rearing experiment (as described above, one plant was randomly selected from each of 45 maternal lines in a common garden). Leaves were taken haphazardly from four different stems, avoiding the youngest and oldest leaves, and combined in a single paper collection envelope; we also avoided damaged leaves, although plants were exposed to constant, low levels of natural herbivory from insect and small mammal herbivores before and during the experiment (thus the present study does not address plasticity of defense in response to herbivore attack). Vacuum-dried, ground leaves (10 mg) were extracted in 2 mL of 70% aqueous ethanol and injected into an Agilent 1200 analytical high performance liquid chromatograph paired with an Agilent 6230 Time-of-Flight mass spectrometer via an electrospray ionization source. Resulting chromatograms were analyzed using MassHunter Quantitative Analysis (v.B.06.00, Agilent, Santa Clara, CA), and major classes of compounds were identified using characteristic relative mass defects (Ekanayaka, Celiz, & Jones, 2015), as described further in the appendix. Leaf protein content was quantified with three replicates (∼2 mg each) per plant using the Bicinchoninic acid assay (Pierce Biotechnology, Waltham, MA). Before grinding, five dried leaflets from each sample were weighed to the nearest 0.1 mg, scanned, and area was measured using ImageJ (v.1.52a); specific leaf area (SLA) was calculated as leaf area divided by dry mass. Finally, leaf toughness was measured on fresh material in the common garden, at the start of the experiment (mid-July, when leaves were also sampled for chemistry and protein) and at the end of the experiment (mid-August), from three leaves per plant at each date, with a penetrometer (Chatillon 516 Series) through the center of the middle leaflet, as in (Harrison et al., 2018); the three leaves were selected haphazardly, avoiding the oldest and youngest leaves. Leaf toughness (averaged across the three leaves per plant at each collection) was correlated between early and late in the season (r = 0.36), but we focus on the measurements taken at the first time point in subsequent analyses for consistency with samples taken at that time for metabolomics.

### 2.3 Overview of analyses of plant traits and caterpillar performance

Our analytical strategy to understand the association between phytochemical variation and caterpillar performance followed two complementary paths, one focusing on reducing the number of variables (through dimension reduction and feature selection) to produce relatively simple models, and the other on the estimation of effects of all individual compounds on caterpillars (without reducing the number of predictor variables). For the first path, involving dimension reduction, we utilized an approach developed for gene transcription studies that identifies groups or modules of correlated variables with hierarchical clustering (Langfelder & Horvath, 2008); after clustering, we reduced the number of independent variables by selecting among modules and other plant traits (specific leaf area, protein and leaf toughness) using lasso regression (Ogutu, Schulz-Streeck, & Piepho, 2012). Lasso regression shrinks coefficients for less important variables to zero, and is thus useful for model selection, in contrast to ridge regression which constrains coefficients (providing stable estimates) while not excluding variables. Modules (and other plant traits) selected in the lasso regression step were subsequently analyzed in Bayesian linear models that are useful in this context because they allowed us to quantify our confidence in the sign of effects (positive or negative) as continuous probabilities (as opposed to relying on arbitrary significance cutoffs). For the second analytical path, we utilized ridge regression (Ogutu et al., 2012) to estimate effects for all compounds simultaneously, which allowed us to investigate the distribution of effects among compounds and classes of compounds. Both analytical paths incorporated cross-validation during the lasso and ridge regressions (further details below in section 2.4.2), and as a means of evaluating the predictive success of the Bayesian models. We also used randomization tests to compare the performance of modules and individual compounds with randomly-chosen suites of compounds.

### 2.4 Dimension reduction and feature selection

#### 2.4.1 Clustering of phytochemical variables

We chose an approach (the first set of analyses mentioned above) that reduces the number of independent variables while allowing us to learn about the correlational structure of the data, specifically unsupervised hierarchical clustering as implemented in the blockwiseModules function of the WGCNA package (Langfelder & Horvath, 2008) in R (R Core Development Team, 2016). Among the options in the pipeline, we used positive correlations among variables (“signed” network type), merge cut height at 0.25, and correlations raised to the power of five (which is where the scale free topology index reached a plateau). Through experimentation, we found that our results with LC-MS data were robust to variation in these choices, including the choice of signed or unsigned networks. After an initial round of clustering, we took a remaining 19 unassigned compounds and put them through a second round of clustering (although the majority of consequential compounds were identified in the first round). One output of the WGCNA procedure is the first eigenvector from each cluster of compounds, which reduced our number of predictor variables by a factor of ten.

#### 2.4.2 Lasso regression and Bayesian models

The resulting eigenvectors plus protein, SLA (specific leaf area) and leaf toughness were then put through the feature reduction step of lasso regression (Ogutu et al., 2012), a penalized regression that allows beta coefficients to be constrained to zero (thus excluding variables). We used the cv.glmnet function of the glmnet package (Friedman, Hastie, Simon, & Tibshirani, 2016) with cross-validation during error reduction set to leave out one plant (and associated caterpillars) at each iteration. The variables selected by the lasso were then put into a Bayesian linear model to estimate coefficients and associated credible intervals using JAGS (version 3.2.0) run in R with the rjags package (Plummer & others, 2003). Two Markov chains were run for 10,000 steps for each analysis (no burn in was required) and chain performance was assessed by plotting chain histories, and calculating the Gelman and Rubin convergence diagnostic and effective sample sizes (Brooks & Gelman, 1998; Gelman, Rubin, & others, 1992). For all models, minimally-influential priors for the regression coefficients were modeled as a normal distribution with a mean of zero and variance of 100 (variance = 1/precision). We quantified our confidence in the sign of coefficients (positive or negative) as the fraction of the posterior samples that were less than zero (for coefficients with a median negative value) or greater than zero (for coefficients with a median positive value).

All analyses were done using the R statistical language (R Core Development Team, 2016) on scaled (z-transformed) predictor variables, and both lasso and Bayesian models used binomial (for survival), Poisson (for development time) and Gaussian (for adult weight) errors. The latter two analyses (development time and adult weight) included sex as a factor. The analysis of development time also included adult weight as a covariate; while (reciprocally) the analysis of adult weight included development time as a predictor. These variables are negatively correlated (r = −0.52), and they function as useful covariates of each other, allowing us to investigate the possibility of unique plant effects on weight gain and development time, which could not be discovered if, for example, these variables were combined into a single performance index.

#### 2.4.3 Cross-validation and resampling to judge model performance

The success of models developed with the dimension reduction and feature selection pipeline was judged in two ways. We used a cross-validation procedure in which we left out one plant (and associated caterpillars) in each iteration of the Bayesian model and then used the estimated coefficients (for phytochemical variables and other plant traits) to predict the performance of the unobserved caterpillars. After 45 iterations (one for each plant), we calculated a simple correlation coefficient between the observed and predicted performance of caterpillars across plants. In addition, we repeatedly resampled the original LC-MS data to match the structure of the reduced set of predictor variables to ask to what extent randomly assembled modules could outperform the empirically-derived modules (in other words, if a model contained two modules with 15 and 20 compounds, simulated predictors would include modules based on 15 and 20 randomly-selected compounds).

### 2.5 Individual compound effects

The second path of our two-part analytical strategy involved simultaneous estimation of the effects of all individual chemical compounds on caterpillar survival, development time and adult weight. For this approach, we again used penalized regression (in the glmnet package [Friedman et al., 2016]), but this time with ridge regression (instead of lasso) which constrains beta coefficients to avoid variance inflation but does not eliminate variables. As with the analyses above, ridge regression was done using error structures appropriate to the specific response variables, and included additional covariates where possible (in models of development time and adult weight). The resulting coefficients associated with all individual compounds were examined as a second perspective on the modules examined in the first set of analyses, and were used to ask to what extent individual compound effects could be predicted by the degree to which they vary among individual plants as quantified with the simple coefficient of variation. To assess confidence in the results of ridge regressions, we used a bootstrap approach, repeatedly resampling the data and estimating coefficients 1000 times, noting the compounds whose bootstrap confidence intervals did or did not overlap zero (Delaney & Chatterjee, 1986). We also allowed for the discovery of interactions among compounds using penalized regression on all individual compounds and all pairwise interactions between compounds. For ease of interpretation, this final analysis of potential interactions used lasso (not ridge) regression, letting the coefficients for many of the individual compounds and pairwise interactions go to zero.

## 3 Results

Of the 450 caterpillars that started the experiment, 261 were reared to eclosion as adults (a mortality rate similar to previous work with this system [Gompert et al., 2015]) on leaves from 45 individual alfalfa plants that were characterized for protein, leaf toughness, specific leaf area and 163 individual metabolomic features (see Figure 1 for variation among plants in caterpillar performance and a subset of plant traits, and appendix Table A1 for a list of compounds). Hierarchical clustering identified 14 subsets (or modules) of compounds with generally low correlations among modules and high correlations within modules (see appendix Figures A1 and A2 for correlations within and among modules, and Figure A3 for module variation among plants). The correlational structure of the phytochemical data is illustrated as an adjacency network in Figure 2 (and in Figure A4 colored by compound class instead of module), where it can be seen that some modules (e.g., modules 1 and 2) contain a great diversity of compound types, while other modules are made up of more narrow classes (e.g. modules 7 and 8 are mostly saponins, a class of defensive metabolites [Levin, 1976]). From the 14 eigenvectors summarizing variation in the modules, as well as the other plant traits, lasso regression (Ogutu et al., 2012) produced a reduced set of potential predictors which were then used in Bayesian multiple regression models that included between six and seven independent variables (Table 1). The models had reasonably high performance in leave-one-out cross-validation: correlations between observed and predicted values were between 0.50 and 0.59 (Table 1), and thus model predictions explained between 25 and 35% of the observed variation in caterpillar performance. Resampling analyses were similarly successful (Supporting Information Figure A5), with only a small fraction (never more than 3%) of randomly-generated models exceeding the variance explained by the models reported in Table 1.

**FIGURE 1.**
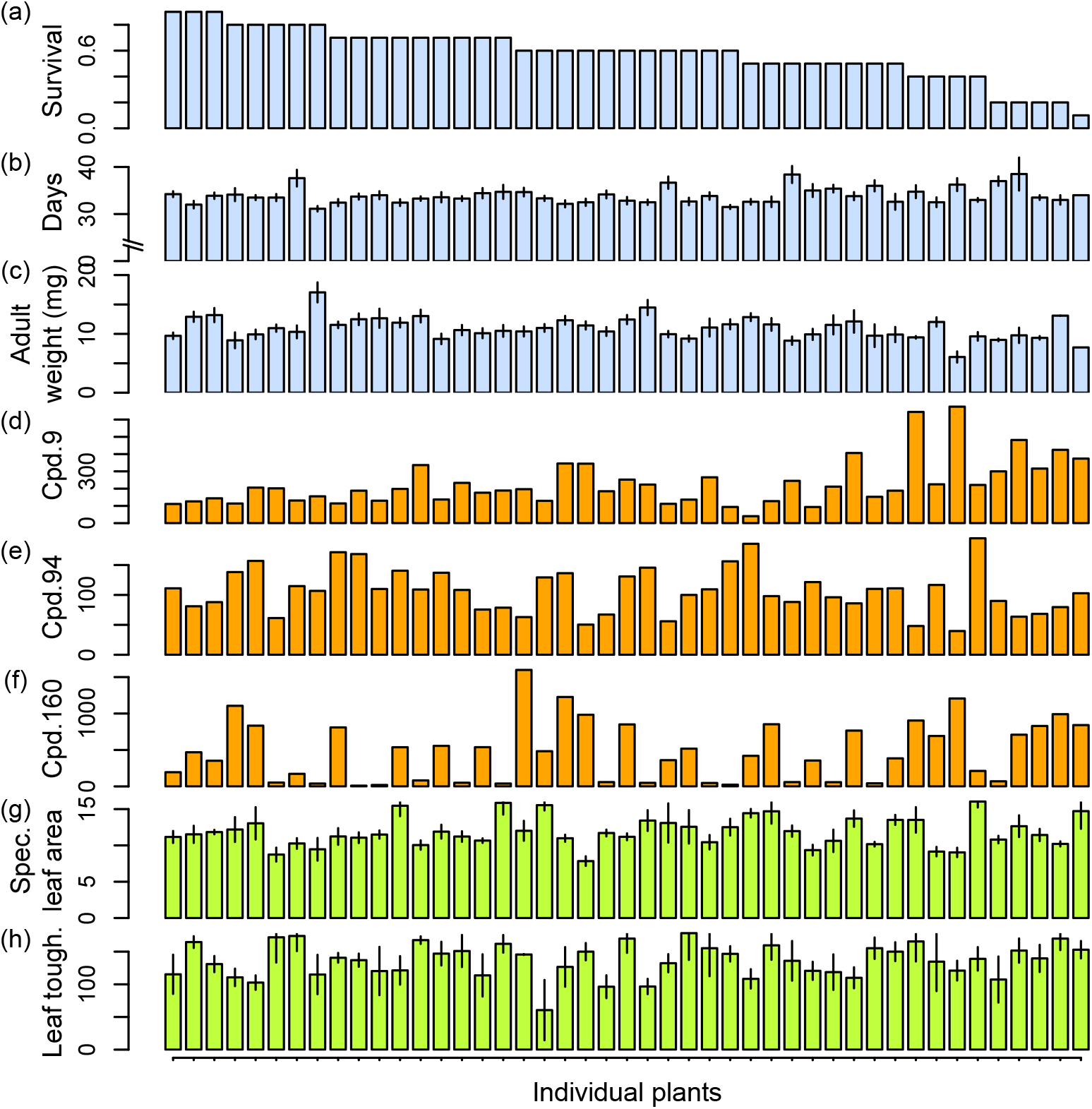
Variation among plants in caterpillar survival (a), development time (b) adult weight (c), three individual compounds (d-e) and two external plant traits, specific leaf area (g) and leaf toughness (h). The three example compounds shown here (out of the 163 assayed) were among the top five most influential compounds for survival, development time and adult weight: cpd. 9 is an alkaloid with a negative association with survival, cpd. 94 (a peptide) has a negative association with development time, and cpd. 160 is a phospholipid with a negative association with adult weight. Individual plants in all panels are organized from left to right by decreasing caterpillar survival in the top panel (a). Standard errors are shown for panels b, c, g and h. The units for d-e are compound relative abundance per dry weight of sample; the units for specific leaf area are cm^2^/mg, and grams/newton for leaf toughness.

**FIGURE 2.**
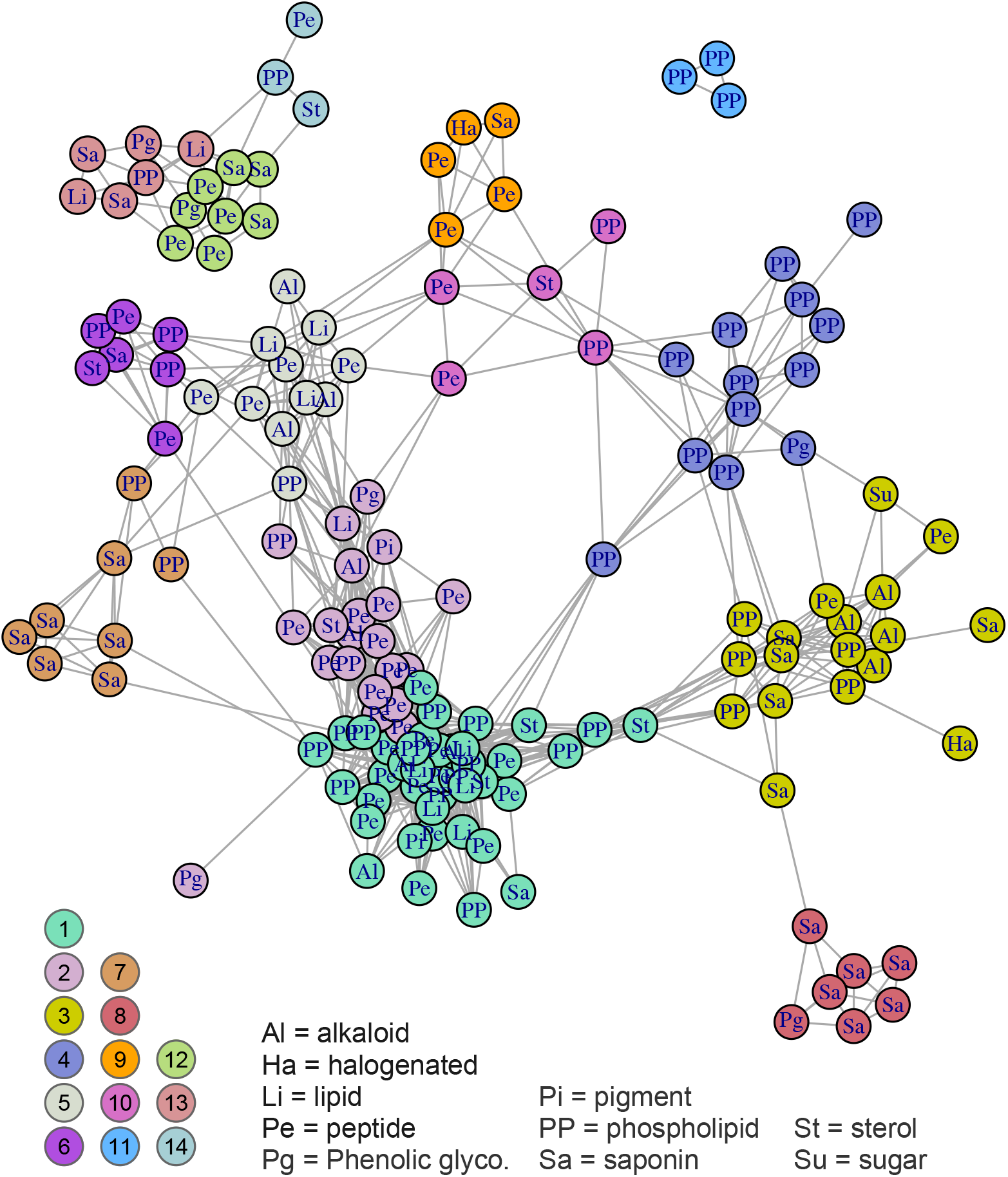
Illustration of correlational structure among compounds: each node in the network is a compound, and compounds are linked by a line if they are correlated among individual plants at 0.5 or above (links among compounds in modules 12-14 represent weaker correlations, greater than 0.1; see main text for details). Nodes are clustered in space for ease of visualization, but relative distances among nodes (and the relative lengths of lines) convey no additional information. Two letter codes within nodes indicate compound classes, as explained in the legend. Colors of nodes correspond to membership in modules as determined by hierarchical cluster analysis; the color key to the 14 modules is shown in the lower left (also see Figure A4 where nodes are colored by compound class). Not shown are a small number of compounds with weak connections to all other compounds, including two compounds that were not included in any module (shown as module zero in Figure 3).

**TABLE 1.** Results from Bayesian regressions of module eigenvectors and covariates predicting caterpillar survival, development time and adult weight (as binomial, Poisson, and Gaussian regressions, respectively, with corresponding units in log-odds, log number of days, and milligrams).

|  | Survival | Development time | Weight |
| --- | --- | --- | --- |
|  | coefficient (CI; prob.) | coefficient (CI, prob.) | coefficient (CI, prob.) |
| m2 | 0.14 (-2.06, 0.48; 0.80) | -0.01 (-0.04, 0.02; 0.77) |  |
| m3 | -0.40 (-0.67, -0.14; >0.99) |  | -0.44 (-0.84, -0.03; 0.98) |
| m4 |  |  | 0.29 (-0.14, 0.70; 0.91) |
| m6 |  | -0.01 (-0.03, 0.01; 0.80) |  |
| m9 | -0.30 (-0.63, 0.03; 0.96) | 0.01 (-0.02, 0.04; 0.79) |  |
| m10 | 0.35 (0.08, 0.62; >0.99) |  | 0.40 (-0.01, 0.82; 0.98) |
| m11 | 0.36 (0.14, 0.58; >0.99) |  |  |
| m13 | -0.18 (-0.43, 0.06; 0.93) |  |  |
| m14 |  | 0.01 (-0.02, 0.03; 0.71) |  |
| SLA | -0.32 (-0.56, -0.08; >0.99) |  | -0.35 (-0.72, 0.012; 0.97) |
| Sex | / | 0.06 (0.02, 0.10; >0.99) | 1.12 (0.40, 1.84; >0.99) |
| mg | / | -0.03 (-0.05, -0.01; >0.99) | / |
| Days | / | / | -1.41 (-1.76, -1.05; >0.99) |
| Intercept | 0.34 (0.14, 0.54; >0.99) | 3.48 (3.45, 3.52; >0.99) | 10.36 (9.81, 10.91; >0.99) |
| Validation | 0.59 | 0.59 | 0.50 |

*Note:* For each regression coefficient, numbers in parentheses are 95% credible intervals (CI, the first two numbers) and the probability that the coefficient has the estimated sign (e.g., 0.80 for the m2 survival coefficient of 0.14 indicates a 80% probability that the m2 module has a positive effect on survival). Note that negative coefficients for development time indicate faster caterpillar development (fewer days) associated with variation in a particular module. Modules (listed in the left column) are only shown if they were included in one of the three regressions following feature selection using lasso regression (see main text for additional details). Empty spaces in the Table appear if a particular module was selected through lasso regression for one or two analyses but not all three (m3, for example, was not selected by lasso regression for development time). Slash marks (/) indicate variables not considered for a particular analysis (e.g., sex, adult weight [mg] and development time [days] were not possible for the survival analysis because they are not observed on dead individuals). Values for “validation” shown in the last row are the correlation between observed and predicted values in cross-validation (Supporting Information Figure S4).

Variation among plants in the suites of covarying compounds had large effects on caterpillar performance: for example, the beta coefficient of −0.40 (on the log-odds scale) associated with module 3 corresponds to a 0.10 reduction in the probability of survival (relative to average) associated with a one unit change in that phytochemical module (Table 1; note that in Table 1 and elsewhere negative coefficients for development time are associated with fewer days, and thus can be thought of as potentially beneficial effects, in contrast to negative coefficients for survival and weight that are detrimental to caterpillars). The phytochemical predictor variables are eigenvectors from clustering analysis, and thus are not entirely straightforward to interpret, especially when the clustering analysis was itself based on z-transformed data. It is important to note that our LC-MS data (used in clustering analysis) consists of peak areas divided by the peak of an internal standard, and again divided by the dry weight of the sample (thus, in total, referred to as “relative abundance per dry weight”; see appendix for additional details including choice of standard). Variation in these numbers reflects variation in concentrations within compounds (among plants), but care should be used in comparing among compounds because of different ionization responses relative to the standard (thus the use of z-transformation for among-compound analyses). Nevertheless, the effects reported in Table 1 reflect real variation in suites of compounds, as can be seen in correlations between eigenvectors and individual compounds in Supporting Information Figure A2, and in variation among plants in average z-scores in Figure A3.

In some cases, modules included in multiple regression models had common effects across response variables (e.g., the positive association of module 10 with both survival and adult weight or the negative association of module 3 also with survival and weight), while other modules had more specific effects on a single response (e.g., modules 11 and 13 on survival). Specific leaf area had a negative association with survival and adult weight, with the coefficients for specific leaf area (−0.32 for survival and −0.35 for weight) being of similar magnitude to some of the phytochemical effects. Neither leaf toughness nor protein had sufficiently strong associations with any of our caterpillar response variables to pass the initial filter of the cross-validated lasso regressions.

Module-based analyses (as in Table 1) focused on feature reduction with lasso regression; as a complementary analytical approach, we also used ridge regression (Ogutu et al., 2012) on all of the individual compounds (ridge regression estimates effects of compounds without excluding variables as in lasso regression). Analyses of individual compounds by ridge regression (Figure 3) were broadly consistent with the strongest module-specific effects, as can be seen, for example, with module 10 having positive effects on survival and adult weight in module analyses (Table 1) and in compound-specific analyses (Figure 3). Similarly, the individual compounds in module 3 had negative compound-specific effects on survival (Figure 3), and that module had the strongest negative effect on survival in the eigenvector-based analyses in Table 1. Not surprisingly, the larger modules (with a greater number of covarying compounds, including many primary metabolites) tended to have a more complex mix of positive and negative effects (for example, modules 1 and 2, Figure 3). For ease of interpretation, the coefficients from compound-specific regressions of survival and development time (in Figures 3 and 4) have been back-transformed to be on the scales of probability and days (respectively), and displayed as changes relative to intercepts. For example, a compound with a relatively large effect on survival in Figure 3 could be associated with a 0.005 reduction in the probability of survival relative to average survival and while holding other compounds constant.

**FIGURE 3.**
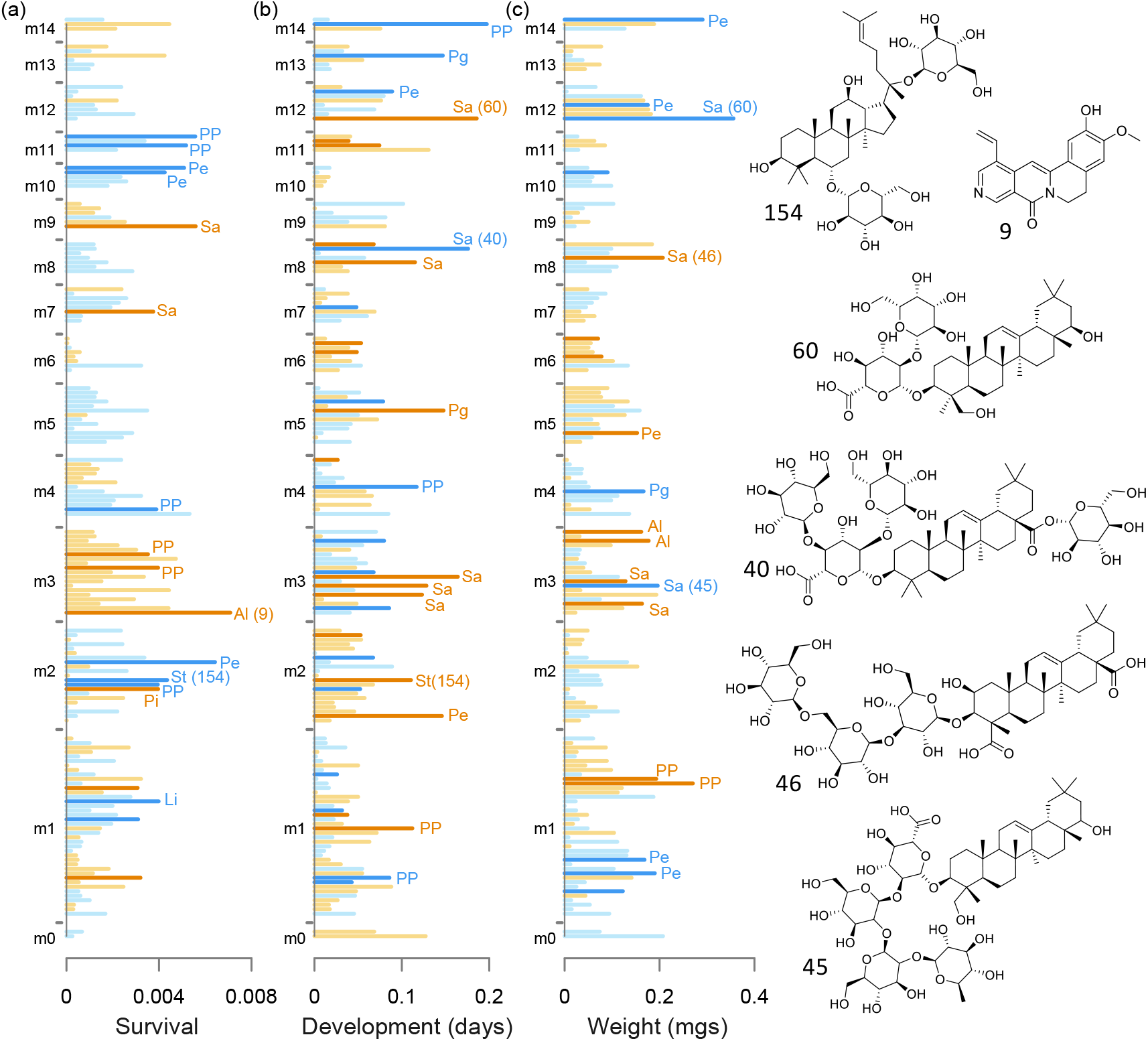
Effects of individual compounds on survival, development time and adult weight, as estimated by ridge regression (using binomial, Poisson and Gaussian models, respectively). The strength of effect for each compound is indicated by the horizontal extent of each bar, and compounds are grouped by modules (m1, m2, etc.); the order of compounds along the vertical axis is arbitrary within modules and fixed across columns. Orange colors indicate negative effects on survival, development and weight, while blue colors are positive effects (note that negative effects for development time correspond to fewer days, or more rapid development). The darker shades of orange and blue mark coefficients whose 95% confidence intervals did not overlap zero in 1,000 bootstrap samples. Values for survival and development time have been back-transformed from units on the log-odds and log scales to units of probability and days to pupation, and are shown as changes from the mean or intercept values. For example, a negative (orange) survival coefficient of 0.005 means a reduction of that amount from the average probability of survival associated with variation in a particular compound. The fifteen compounds with the largest coefficients (by absolute value) and bootstrap intervals not overlapping zero are labelled by compound classes (see Figure 2 for abbreviations) in each panel. Structural annotations are shown to the right for six compounds based on matches from the METLIN metabolomics database, as follows by compound number: 154 (unidentified sterol); 9 (unidentified alkaloid); 60 (soyasaponin A3); 40 (unidentified saponin); 46 (medicagenic acid 3-O-triglucoside); 45 (medinoside E). Those same compounds are identified in parentheses in the main panels next to the bars corresponding to their individual effects.

**FIGURE 4.**
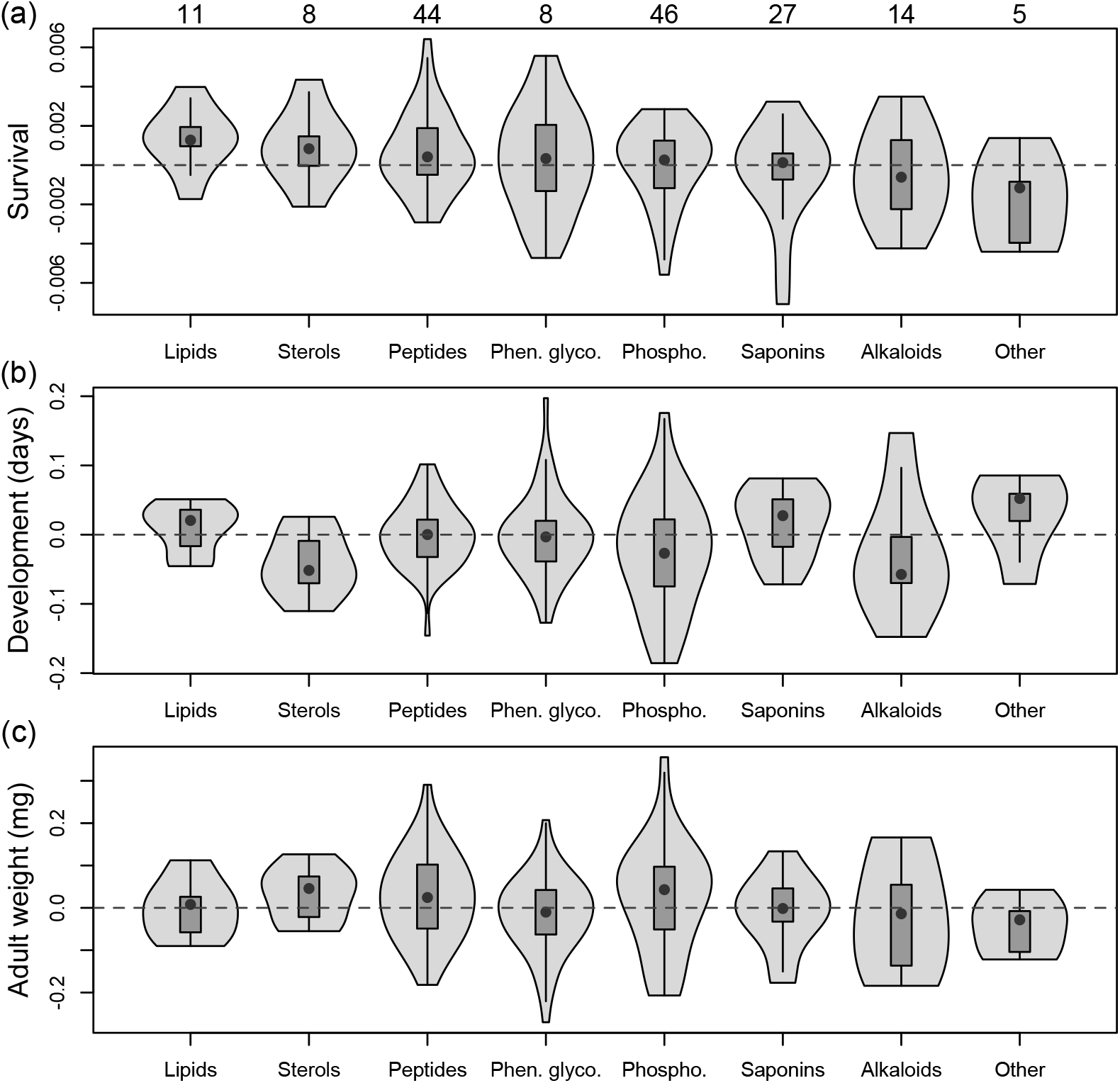
Violin plots of compound-specific effects (coefficients from ridge regressions) summarized by chemical classes. Sample sizes for each category are shown above the top panel (“Other” includes 1 sugar, 2 pigments and 2 halogenated compounds). Categories are arranged from left to right based on the gradient of median positive to negative effects on survival. Coefficients for survival (a) and development time (b) have been back-transformed from the units of log-odds and log to probability and days to pupation, respectively, and shown as deviations from the mean or intercept value (as in Figure 3). Note that negative effects for development time correspond to fewer days (more rapid development). Violin plots show medians (black dots) and interquartile ranges (boxes); vertical lines are upper and lower fences (the third quartile + 1.5 * the interquartile range, and the lower quartile - 1.5 * the interquartile range, respectively) surrounded by kernel density envelopes.

We saw some variation among classes of compounds in their effects on caterpillars (Figure 4). All classes included positive and negative effects, with saponins, alkaloids and phenolic glycosides including some of the stronger negative effects of individual compounds, while lipids and sterols tended towards positive associations with survival and development (Figure 4). We also considered potential pairwise interactions among individual compounds, and found few interactions that passed the filter of the penalized regression (Supporting Information Table A2), at least relative to the large number of potential interactions. Saponins and alkaloids tended to be overrepresented in the interactions that were detected, and phenolic glycosides were involved in stronger negative interactions relative to other compounds (Figure A6). We did not find evidence that more or less variable compounds (among individual plants) had differential effects on caterpillars (Figure A7).

## 4 Discussion

The results reported here represent a dissection of the phytochemical landscape facing a specialized insect herbivore attacking a recently-colonized host plant (Hunter, 2016). The phytochemical landscape is both physical, referring to variation in compounds among individual plants in a common garden (Figure A3), and hypothetical to the extent that effects of individual compounds on caterpillars are estimated, although compounds are of course not encountered in isolation. Our exploration of the phytochemical landscape facing *L. melissa* on *M. sativa* is necessarily a first draft based on a single point in time. Despite the snapshot nature of our study, models including suites of covarying metabolites and other plant traits had predictive success for caterpillar performance and suggested different natural products affecting survival, development time and adult weight. Previous work with *M. sativa* and other insect herbivores has focused on sapogenic glycosides (Levin, 1976), and a simple outcome from our study could have been that one or a small number of saponins have anti-herbivore properties that reduce fitness of our focal insect. Instead, we find large numbers of compounds with potentially consequential effects on caterpillars (Figure 3), and which were in some cases of similar magnitude or greater than the effects of morphological features, including leaf toughness and specific leaf area (Carmona, Lajeunesse, & Johnson, 2011).

We find that prominent classes of specialized metabolites in our focal plants, such as saponins and peptides, include compounds with both positive and negative effects on survival and development of caterpillars. Positive effects of these compounds are potentially associated with feeding stimulation, as has been observed (along with other positive effects) for other specialist herbivores and plant toxins (Seigler & Price, 1976; Smilanich, Fincher, & Dyer, 2016). Negative effects of saponins on insects potentially include disruption of hormone production (Chaieb, 2010), although exact modes of action on *L. melissa* will await further study. Although many of the compounds with strong effects are specialized metabolites (including alkaloids and phenolic glycosides, as well as saponins and peptides), we also find both positive and negative effects on caterpillar performance associated with primary metabolites (Figure 4), especially phospholipids (Figure 2). These could be direct effects if a compound is suboptimal for development, or they could be associated with nutritional imbalance (Behmer, 2009), such that too much of one nutrient makes it difficult for caterpillars to consume a balanced diet. It has been suggested that the presentation of unbalanced nutrition can be a kind of anti-herbivore strategy (Berenbaum, 1995), although this has not been studied in the *L. melissa*-*M. sativa* interaction.

The finding that our specialist herbivore is affected by a wide range of metabolites that vary greatly even within a single host population has implications for our understanding of heterogeneity in the system, and also for local adaptation of the herbivore to the novel host. *Lycaeides melissa* typically colonizes weedy or feral patches of *M. sativa* on roadsides or integrated into natural communities, and previous work has documented dramatic variation among individual alfalfa locations (often in close proximity) in the extent to which they can support caterpillar development (Harrison et al., 2016). Previous phytochemical data with a lower resolution was less successful in explaining that variation (Harrison et al., 2016), but the results reported here suggest that among-patch variation could be explained by future studies using detailed metabolomic data. The within-population complexity described in the current study also raises the possibility that the novel host presents a multi-faceted and potentially ever-shifting target from the perspective of evolving butterfly populations (Chaturvedi et al., 2018; Gompert et al., 2015; Harrison et al., 2016). In particular, it is possible that *M. sativa* defense against a specialist herbivore might be realized through different combinations (within and among populations) of individually-acting compounds, thus making it less likely that butterflies in any one population possess an effective suite of alleles that improve fitness on *M. sativa*.

The correlational structure of the phytochemical variation that we observed has implications for the evolution of plant defense and the accumulation of insect herbivores on *M. sativa*. Specifically, correlations among modules (which are themselves composed of a diversity of compound types) should make it possible to hypothesize directions of least resistance for defense evolution. Module 3, including an alkaloid with a prominent effect on caterpillars but also phospholipids and saponins, had a negative effect on survival (Table 1, Figure 3). Module 3 negatively covaried with module 2, which was itself positively associated with caterpillar survival (including a peptide of large effect but many other compound types as well). Thus an increase in module 3 and an associated decrease in 2 would be beneficial for the plant, at least with respect to herbivory by our focal herbivore. Predicting evolutionary response by *M. sativa* would of course depend on a genetic understanding of the relevant plant traits, which the present study does not include. However, a recent study of *M. melissa* performance on a related plant, *Medicago truncatula*, found that genetic variation in the plant explained a substantial proportion of phenotypic variation (between 8 and 57%) in phytochemical and structural traits but also in caterpillar performance (Gompert et al., 2019). Of course, most plants do not have the luxury of optimizing defense against a single herbivore, and it is easy to imagine that improvements in defense against one enemy could lead to increased attraction to another (Salazar et al., 2018), especially given the diversity of effects even within major classes studied here, including saponins and phenolic glycosides. Compounds in the latter class (phenolics) were found to have strong positive and negative effects on assemblages of arthropods associated with the maternal plants from which seeds were collected to start the common garden used in the present study (Harrison et al., 2018).

The results reported here raise a number of avenues for future exploration, including the apparent overrepresentation of both saponins and alkaloids in interactions with other compounds (Figure A6). Relative mass defect (RMD) is a useful tool for the categorization of compounds (Table A1), but it has limitations in complex mixtures; we are developing methods that use other data from high resolution mass spectrometry to further refine categorization of *Medicago* metabolites (Philbin & Forister, *in prep*). Also, in the present study, we have not attempted to separate constitutive and induced defenses (Jansen et al., 2009) as the plants in the common garden were exposed to natural and continuous levels of herbivory. We also acknowledge that feeding under laboratory conditions is of course not natural, although we found in a previous study that genetic variants (in caterpillars) associated with success in laboratory feeding trials were at least partially predictive of genetic variation associated with alfalfa use by *L. melissa* in the wild (Chaturvedi et al., 2018). Thus it is clear that metabolomic data, such as those analyzed here, have the potential to both open up new avenues of conceptual development in plant-insect interactions and to link micro-evolutionary trajectories across hosts and herbivores.

## ACKNOWLEDGEMENTS

This work was supported by National Science Foundation grant DEB-1638793 to MLF and CDD, DEB-1638768 to ZG, DEB-1638773 to CCN, DEB-1638922 to JAF, and DEB-1638602 to CAB; MLF was additionally supported by a Trevor James McMinn professorship. Thanks to Ian Wallace, the Hitchcock Center for Chemical Ecology and the PIG group at UNR for discussion and expertise.

## AUTHOR CONTRIBUTIONS

MLF: designed experiment, conducted analyses, wrote first draft. SY: conducted experiment and contributed to experimental design. CSP, CDD, BH: generated and interpreted phytochemistry and protein data. MLF, JGH, OS: developed and maintained common garden. JAF, ZHM, CCN, LAR: contributed to analyses and experimental design. CAB, JAF, ZG, CCN: contributed to experimental design. All authors: contributed to writing.

## DATA AVAILABILITY STATEMENT

Data will be made available from the Dryad Digital Repository: *link to be determined*.

## APPENDIX

### Additional phytochemical methods 1: LC-TOF analysis of foliar plant tissue

Foliar tissue was dried *in vacuo* and individual leaves were selected haphazardly from individual plants and finely ground (TissueLyser II, Quiagen; Hilden, Germany). Approximately 10 mg of ground foliar tissue was weighed and extracted in 2.00 ml of 70% aqueous ethanol, and briefly vortexed before 15 minutes of sonication. This suspension was centrifuged (500 rpm) for 10 min, then 1 ml aliquots of the supernatant were filtered through a 96-well filter (AcroPrep, 1 mL, 1 μm glass fiber) into glass vial inserts and capped with a silicone mat before analysis. Chromatography was performed on an Agilent 1200 analytical HPLC equipped with a binary pump, autosampler, column compartment and diode array UV detector, coupled to an Agilent 6230 Time-of-Flight mass spectrometer via an electrospray ionization source (ESI-TOF; gas temperature: 325 °C, flow: 10 L/m; nebulizer pressure: 35 psig; VCap: 3500 V; fragmentor: 165 V; skimmer: 65 V; octopole: 750 V). Extracts (1.00 μL) were co-injected with 0.50 μL of digitoxin internal standard (0.200 mM, Sigma-Aldrich) and eluted at 0.500 mL/min through a Kinetex EVO C18 column (Phenomenex, 2.1 x 100 mm, 2.6 μ, 100 Å) at 40 °C. The linear binary gradient was comprised of buffers A (water containing 0.1 % formic acid) and B (acetonitrile containing 0.1 % formic acid) changing over 30 minutes accordingly: 0-1 min 5% B, ramp to 50% B at 4 min, ramp to 100% B at 21 min, 21-25 min 100% B ramping to 1.00 mL/min, before re-equilibrating the column from 25-30 min at 5% B, 0.5 mL/min.

Individual compounds were quantified relative to the digitoxin internal standard using Agilent MassHunter Quantitative Analysis. Digitoxin is a commercially available cardenolide which has previously been used as an internal standard in saponin analysis (Balsevich, Bishop, & Deibert, 2009). Its structural similarity to, and lack of coelution with, saponins and its absence in *Medicago* extracts make it an ideal internal standard for quantitation of saponins, the focal phytochemical class in this study. While digitoxin allows for the quantitation of saponins as “digitoxin equivalents”, this does not extend to other phytochemical classes due to differences in ionization efficiency inherent in structural differences for other classes. In these cases, the digitoxin internal standard still serves to partially correct for between run variation in instrument response. We do not make quantitative assertions between phytochemical classes, only assertions based on their within-class variation for this reason.

Putative phenolics (200-400 ppm) and saponins (400-650 ppm) were annotated using the relative mass defect (RMD) characteristic of each phytochemical (see next appendix section). Compounds with RMD greater than 650 were presumed to be lipids or sterols. These assignments were revised by identifying presumed peptides based on even *m/z* features. Mass spectra of presumed phenolics, saponins and lipids were cross-referenced against the METLIN database (Smith et al., 2005) to further categorize annotations into phospholipids, vitamins (vitamin D derivatives), carotenoids, sterols, amino acids, alkaloids and sugars. One compound displayed an isotope distribution characteristic of a chlorinated structure and was designated as being halogenated. Due to the lack of structural information in ESI-TOF mass spectra, annotation beyond this classification was not possible.

### Additional phytochemical methods 2: relative mass defect (RMD)

Relative mass defect is a recently-developed method for inferring structural information from high resolution mass spectrometry data (Ekanayaka et al., 2015) which we have used to aid in the classification of metabolites in *M. sativa* and to a lesser extent propose putative structures. Here we describe the theoretical background for the calculation and use of relative mass defect. Mass defect is the deviation of atomic mass (see definitions below) from its mass number (e.g. hydrogen: *A_H_* = 1, *m_H_* = 1.00784 Da, *d_H_* = 0.00784). Relative mass defect of an atom (RMD_a_) in ppm is calculated as:

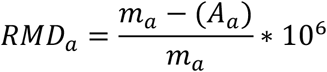

Where *m_a_* is the atomic mass and *A_a_* is the mass number of that atom. Although H has a positive mass defect (RMD_H_ = 7740ppm), mass loss due to the strong nuclear force (Einstein, 1905) leads to increasingly negative mass defect as *A* increases. Atoms commonly found in natural products (RMD_N_ = 221, RMD_C_ = 0, RMD_O_ = −319, RMD_P_ = −846, RMD_S_ = −873 ppm) have a relative mass defect which is a full order of magnitude lower than that of hydrogen. As a result, the relative mass defect (RMD_M_) of a molecule estimates the number of hydrogen atoms relative to other atoms in a natural product: a high RMD molecule has a higher %H than a low RMD molecule. When chemical formula is known, computing the theoretical RMD_M_ of a molecule M is facile:

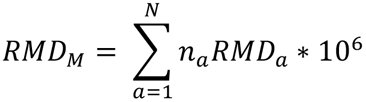

Where *n_a_* is the count of the *a*th element in the set of elements composing a natural product, and *N* is the number of elements in that natural product. As a chemical formula this would appear in the form C_nC_H_nH_N_nN_O_nO_P_nP_S_nS_. When the chemical formula of a natural product is unknown, the RMD of a molecule M can be calculated from HRMS data:

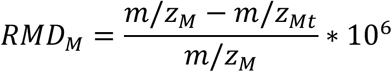

Where *m/z_M_* and *m/z_Mt_* are the mass to charge ratio and nominal mass-to-charge ratio of molecular ion M, respectively.

The RMD of a molecular ion then serves as an experimental estimate of degree of unsaturation; or more generally, how many H are present per unit of molecular mass. This metric can be very useful for discriminating natural products which may have the same nominal mass but differ in exact mass, obtained via HRMS data which is accurate at a sub-ppm level. In Figure A1 (left), three structures are listed which have the same nominal mass (270 Da) but have increasing RMD as the contribution of H to the exact mass increases. Pinostrobin, the most unsaturated and oxidized molecule has the lowest %H and lowest RMD (330 ppm) and as the %H increases to estrone and then methyl palmitate, the RMD of each molecule increases (600 ppm and 948 ppm respectively). This approach is useful when trying to discriminate flavonoid glycosides, such as the apigenin glycoside (Figure A8, right, RMD = 255 ppm) from saponins such as medicagenic acid glycoside (Figure A8, right, RMD = 494) in *M. sativa* extracts.

However, other phytochemicals in these extracts may also have similar RMD, and this metric should not be solely relied upon for annotation. Daughter ions may also differ in RMD from their parent ions due to fragmentation or loss of H to form cationic species. Other information, such as relative retention time, molecular mass and odd molecular masses indicating nitrogenous compounds can also inform classification and annotation.

Definitions:

**Atomic mass (*m*):** exact mass of an atom measured in daltons (Da).

**Mass number (*A*):** the total number of nucleons (protons + neutrons) in an atom.

**Mass to charge ratio (*m/z*):** ion mass divided by charge. The measured unit of mass in a mass spectrometer. When *z* = 1, *m/z* equals ion mass.

**Nominal mass to charge raio (*m/z_t_*):** mass to charge ratio truncated to zero decimals (or floor function). For example, the nominal mass of *m/z =* 270.2559 is *m/z_t_* = 270.

**High Resolution Mass Spectrometry (HRMS):** Mass spectrometric techniques which yield masses accurate to four decimal places which allows for prediction of putative chemical formulae.

**Isobaric:** molecules having the same mass

**Molecular ion:** Ionic species representing an intact molecule

**Parent ion:** Molecular ion which becomes fragmented into daughter ions

**Daughter ions:** Resultant ions from the fragmentation of parent ions

---

**TABLE A1.** Details on individual compounds (numbered along the left column), including module numbers (see Figure 1), chemical class assignments, ridge regression beta coefficients (for caterpillar survival, development time and adult weight as in Figure 3), and mass spectra results: *m/z*, retention time (RT ave.) and relative mass defect

| Cpd. | Module | Class | Surv. | Dev. | Weight | $m/z$ | RT ave. | RMD |
| --- | --- | --- | --- | --- | --- | --- | --- | --- |
| 65 | m. 0 | Phospholipid (PP) | 0.00026 | -0.1273 | 0.2072 | 299.2719 | 6.451733 | 909 |
| 180 | m. 0 | Sterol (St) | 0.00067 | -0.0684 | 0.0740 | 485.3591 | 27.309711 | 740 |
| 23 | m. 1 | Peptide (Pe) | 0.00065 | -0.0629 | 0.1109 | 492.2455 | 4.985533 | 499 |
| 30 | m. 1 | Alkaloid (Al) | 0.00059 | 0.0172 | -0.0105 | 366.1097 | 5.301200 | 300 |
| 35 | m. 1 | Saponin (Sa) | 0.00020 | 0.0112 | 0.1319 | 459.2233 | 5.533244 | 486 |
| 37 | m. 1 | Peptide (Pe) | -0.00041 | 0.0073 | 0.1304 | 1564.7065 | 5.623800 | 452 |
| 50 | m. 1 | Peptide (Pe) | -0.00048 | -0.0165 | 0.1683 | 1238.5847 | 5.966289 | 472 |
| 53 | m. 1 | Peptide (Pe) | -0.00042 | -0.0301 | 0.0121 | 1106.5421 | 6.010844 | 490 |
| 57 | m. 1 | Peptide (Pe) | -0.00183 | 0.0547 | 0.1034 | 256.2642 | 6.141222 | 1031 |
| 58 | m. 1 | Peptide (Pe) | -0.00114 | -0.0546 | 0.1899 | 1192.5417 | 6.211267 | 454 |
| 63 | m. 1 | Phospholipid (PP) | -0.00320 | 0.0857 | -0.1410 | 518.3253 | 6.374000 | 628 |
| 64 | m. 1 | Peptide (Pe) | -0.00053 | 0.0427 | 0.0425 | 562.322 | 6.399089 | 573 |
| 67 | m. 1 | Phospholipid (PP) | -0.00246 | -0.0880 | 0.0248 | 284.2959 | 6.617089 | 1041 |
| 70 | m. 1 | Peptide (Pe) | 0.00052 | -0.0476 | 0.1226 | 1168.6682 | 7.022178 | 572 |
| 78 | m. 1 | Peptide (Pe) | 0.00060 | 0.0464 | -0.0439 | 694.402 | 7.380578 | 579 |
| 84 | m. 1 | Phospholipid (PP) | 0.00100 | -0.0265 | 0.0137 | 293.2112 | 8.701933 | 720 |
| 89 | m. 1 | Phospholipid (PP) | -0.00032 | -0.0165 | 0.0532 | 546.3438 | 10.332689 | 629 |
| 93 | m. 1 | Alkaloid (Al) | -0.00029 | -0.0176 | 0.0149 | 358.2468 | 10.897978 | 689 |
| 95 | m. 1 | Lipid (Li) | 0.00168 | 0.0453 | 0.0941 | 425.2903 | 11.479511 | 683 |
| 111 | m. 1 | Alkaloid (Al) | -0.00022 | 0.0119 | 0.0602 | 592.5035 | 15.707178 | 850 |
| 113 | m. 1 | Sterol (St) | 0.00100 | 0.0130 | -0.0143 | 661.3032 | 15.991800 | 458 |
| 128 | m. 1 | Peptide (Pe) | -0.00269 | 0.0354 | -0.0869 | 584.4235 | 17.597156 | 725 |
| 129 | m. 1 | Peptide (Pe) | -0.00106 | -0.0033 | -0.0254 | 567.4209 | 17.818844 | 742 |
| 136 | m. 1 | Peptide (Pe) | 0.00051 | 0.0017 | 0.0215 | 932.6342 | 18.698356 | 680 |
| 139 | m. 1 | Phospholipid (PP) | 0.00205 | 0.0077 | -0.0888 | 792.5648 | 18.853422 | 713 |
| 146 | m. 1 | Sterol (St) | -0.00004 | -0.0499 | -0.0437 | 451.3596 | 20.131378 | 797 |
| 149 | m. 1 | Phospholipid (PP) | -0.00047 | 0.0087 | -0.0978 | 784.5599 | 20.769978 | 714 |
| 157 | m. 1 | Sterol (St) | 0.00117 | 0.0258 | 0.0326 | 429.3741 | 21.812222 | 871 |
| 160 | m. 1 | Phospholipid (PP) | -0.00322 | 0.0023 | -0.1929 | 747.6095 | 21.920444 | 815 |
| 161 | m. 1 | Phospholipid (PP) | 0.00061 | 0.0140 | -0.2701 | 552.4296 | 22.245667 | 778 |
| 162 | m. 1 | Phospholipid (PP) | -0.00307 | 0.0165 | -0.1205 | 907.5367 | 22.382756 | 591 |
| 163 | m. 1 | Peptide (Pe) | -0.00153 | -0.0016 | -0.1119 | 886.5581 | 22.381711 | 630 |
| 164 | m. 1 | Peptide (Pe) | 0.00277 | -0.0499 | 0.1871 | 830.676 | 22.432867 | 814 |
| 165 | m. 1 | Lipid (Li) | 0.00398 | -0.0385 | 0.0233 | 537.5365 | 22.754333 | 998 |
| 167 | m. 1 | Phospholipid (PP) | 0.00199 | 0.0207 | 0.0001 | 708.5134 | 23.041267 | 725 |
| 169 | m. 1 | Lipid (Li) | 0.00100 | 0.0316 | 0.0242 | 565.5681 | 23.807289 | 1004 |
| 170 | m. 1 | Phospholipid (PP) | 0.00214 | -0.0376 | -0.0473 | 885.5552 | 24.037489 | 627 |
| 171 | m. 1 | Lipid (Li) | 0.00310 | 0.0220 | 0.0281 | 536.4387 | 24.087178 | 818 |
| 172 | m. 1 | Phospholipid (PP) | 0.00194 | -0.0314 | -0.0176 | 869.559 | 24.216778 | 643 |
| 173 | m. 1 | Phospholipid (PP) | -0.00145 | -0.1120 | 0.0485 | 787.427 | 24.262511 | 542 |
| 175 | m. 1 | Pigment (Pi) | 0.00138 | -0.0712 | -0.1041 | 871.5766 | 24.897822 | 662 |
| 177 | m. 1 | Lipid (Li) | -0.00052 | 0.0206 | 0.0080 | 208.1347 | 27.036422 | 647 |
| 18 | m. 2 | Phen. glyco. (Pg) | -0.00246 | -0.0574 | 0.0201 | 323.0738 | 4.163533 | 228 |
| 33 | m. 2 | Phen. glyco. (Pg) | -0.00040 | -0.0210 | -0.0409 | 493.1329 | 5.430044 | 505 |
| 69 | m. 2 | Peptide (Pe) | 0.00001 | -0.0226 | -0.0657 | 1018.537 | 6.846578 | 527 |
| 92 | m. 2 | Lipid (Li) | 0.00220 | -0.0455 | 0.1122 | 375.2549 | 10.640356 | 679 |
| 94 | m. 2 | Peptide (Pe) | 0.00041 | -0.1458 | 0.0496 | 618.4291 | 11.238667 | 694 |
| 99 | m. 2 | Alkaloid (Al) | -0.00001 | -0.0174 | -0.0306 | 618.4291 | 12.470689 | 694 |
| 100 | m. 2 | Peptide (Pe) | 0.00235 | -0.0290 | -0.0485 | 600.4193 | 13.376289 | 698 |
| 101 | m. 2 | Alkaloid (Al) | 0.00040 | -0.0528 | 0.0074 | 618.4291 | 13.049600 | 694 |
| 103 | m. 2 | Peptide (Pe) | -0.00012 | -0.0534 | -0.0377 | 603.4311 | 13.358222 | 714 |
| 104 | m. 2 | Peptide (Pe) | 0.00242 | -0.0386 | -0.0329 | 600.4193 | 13.376289 | 698 |
| 109 | m. 2 | Peptide (Pe) | 0.00025 | -0.0441 | 0.0031 | 600.4193 | 15.398533 | 698 |
| 112 | m. 2 | Peptide (Pe) | -0.00036 | 0.0008 | -0.0056 | 600.4193 | 15.860356 | 698 |
| 114 | m. 2 | Peptide (Pe) | 0.00337 | 0.0671 | 0.0466 | 584.4235 | 16.288622 | 725 |
| 122 | m. 2 | Peptide (Pe) | 0.00642 | 0.0166 | 0.1315 | 584.4235 | 16.758622 | 725 |
| 133 | m. 2 | Phospholipid (PP) | -0.00094 | 0.0888 | -0.1533 | 716.5243 | 18.284733 | 732 |
| 134 | m. 2 | Peptide (Pe) | 0.00260 | 0.0043 | 0.0275 | 568.4294 | 18.417311 | 755 |
| 135 | m. 2 | Peptide (Pe) | -0.00007 | -0.0027 | 0.0696 | 937.5896 | 18.675222 | 629 |
| 154 | m. 2 | Sterol (St) | 0.00435 | -0.1104 | 0.0743 | 693.425 | 21.485978 | 613 |
| 155 | m. 2 | Phospholipid (PP) | 0.00395 | -0.0670 | 0.0774 | 635.3816 | 21.488511 | 601 |
| 166 | m. 2 | Pigment (Pi) | -0.00396 | 0.0522 | -0.0076 | 871.5766 | 22.912311 | 662 |
| 168 | m. 2 | Peptide (Pe) | 0.00091 | -0.0479 | 0.0062 | 1057.7337 | 23.685733 | 694 |
| 2 | m. 3 | Halogen (Ha) | -0.00085 | 0.0590 | 0.0426 | 199.0946 | 0.546622 | 475 |
| 5 | m. 3 | Alkaloid (Al) | -0.00443 | 0.0446 | -0.0332 | 217.0692 | 0.628311 | 485 |
| 6 | m. 3 | Peptide (Pe) | -0.00292 | -0.0089 | 0.0744 | 309.1795 | 0.638667 | 581 |
| 7 | m. 3 | Sugar (Su) | -0.00441 | 0.0855 | -0.1219 | 365.1066 | 0.653400 | 292 |
| 9 | m. 3 | Alkaloid (Al) | -0.00708 | 0.0404 | -0.0229 | 321.1156 | 0.696467 | 360 |
| 12 | m. 3 | Alkaloid (Al) | -0.00114 | 0.0704 | -0.1614 | 186.0765 | 0.831267 | 411 |
| 13 | m. 3 | Alkaloid (Al) | -0.00088 | 0.0796 | -0.1770 | 140.071 | 0.843044 | 1008 |
| 14 | m. 3 | Peptide (Pe) | -0.00221 | 0.0548 | -0.0971 | 225.0855 | 1.036511 | 380 |
| 34 | m. 3 | Phospholipid (PP) | -0.00396 | -0.0466 | -0.0396 | 547.3472 | 5.507867 | 634 |
| 38 | m. 3 | Saponin (Sa) | -0.00193 | 0.0677 | -0.0541 | 633.348 | 5.635956 | 549 |
| 41 | m. 3 | Saponin (Sa) | -0.00335 | -0.1640 | 0.1125 | 471.3488 | 5.634111 | 740 |
| 42 | m. 3 | Saponin (Sa) | -0.00151 | 0.0292 | -0.1282 | 631.3333 | 5.725444 | 816 |
| 45 | m. 3 | Saponin (Sa) | -0.00020 | -0.1281 | 0.1960 | 1105.5828 | 5.781400 | 527 |
| 55 | m. 3 | Saponin (Sa) | -0.00096 | -0.1231 | -0.1934 | 617.3527 | 6.043333 | 678 |
| 62 | m. 3 | Saponin (Sa) | -0.00139 | -0.0482 | -0.1629 | 825.4299 | 6.314022 | 521 |
| 120 | m. 3 | Phospholipid (PP) | -0.00122 | -0.0073 | -0.0314 | 732.5644 | 16.529933 | 770 |
| 141 | m. 3 | Phospholipid (PP) | -0.00302 | -0.0398 | 0.0321 | 963.6051 | 18.994089 | 628 |
| 143 | m. 3 | Phospholipid (PP) | -0.00352 | 0.0180 | 0.0291 | 951.6048 | 19.629644 | 636 |
| 174 | m. 3 | Phospholipid (PP) | -0.00473 | 0.0474 | -0.0259 | 805.6846 | 23.370044 | 759 |
| 27 | m. 4 | Phen. glyco. (Pg) | 0.00157 | -0.0575 | 0.1663 | 441.2984 | 5.217978 | 676 |
| 80 | m. 4 | Phospholipid (PP) | 0.00323 | -0.0657 | 0.1113 | 518.3253 | 7.756911 | 628 |
| 83 | m. 4 | Phospholipid (PP) | 0.00205 | 0.0001 | 0.0979 | 520.3405 | 8.404400 | 654 |
| 85 | m. 4 | Phospholipid (PP) | 0.00189 | -0.0632 | -0.0102 | 454.2938 | 8.763000 | 647 |
| 86 | m. 4 | Phospholipid (PP) | 0.00389 | -0.0043 | -0.0528 | 496.3412 | 8.889378 | 687 |
| 91 | m. 4 | Phospholipid (PP) | 0.00532 | 0.0847 | 0.1361 | 524.3727 | 10.499178 | 711 |
| 140 | m. 4 | Phospholipid (PP) | 0.00237 | -0.0266 | -0.0049 | 797.5204 | 18.879244 | 653 |
| 142 | m. 4 | Phospholipid (PP) | -0.00099 | 0.0177 | 0.0110 | 714.5088 | 19.293533 | 711 |
| 144 | m. 4 | Phospholipid (PP) | -0.00135 | 0.0014 | 0.0366 | 756.5562 | 19.707467 | 735 |
| 148 | m. 4 | Phospholipid (PP) | -0.00123 | 0.0070 | 0.0349 | 716.525 | 20.684533 | 732 |
| 151 | m. 4 | Phospholipid (PP) | -0.00066 | 0.0324 | -0.0100 | 801.5511 | 20.976333 | 688 |
| 153 | m. 4 | Phospholipid (PP) | -0.00212 | 0.0224 | 0.0440 | 758.5721 | 21.488156 | 754 |
| 159 | m. 4 | Phospholipid (PP) | 0.00041 | 0.1169 | 0.0507 | 803.5978 | 22.069778 | 744 |
| 4 | m. 5 | Alkaloid (Al) | 0.00060 | -0.0718 | 0.0563 | 104.1077 | 0.595533 | 1035 |
| 8 | m. 5 | Alkaloid (Al) | 0.00026 | 0.0377 | 0.0717 | 144.1024 | 0.663689 | 711 |
| 10 | m. 5 | Lipid (Li) | 0.00096 | 0.0048 | -0.0902 | 301.2133 | 0.700111 | 708 |
| 11 | m. 5 | Alkaloid (Al) | 0.00122 | -0.0359 | -0.0768 | 158.1181 | 0.705733 | 747 |
| 15 | m. 5 | Peptide (Pe) | 0.00110 | -0.0136 | 0.1019 | 188.0714 | 2.198778 | 380 |
| 22 | m. 5 | Phen. glyco. (Pg) | 0.00349 | -0.1477 | 0.1581 | 623.1272 | 5.055600 | 506 |
| 32 | m. 5 | Phen. glyco. (Pg) | -0.00083 | 0.0498 | -0.1267 | 507.1149 | 5.435600 | 638 |
| 54 | m. 5 | Peptide (Pe) | 0.00129 | 0.0416 | -0.0698 | 351.2153 | 6.034822 | 613 |
| 87 | m. 5 | Peptide (Pe) | 0.00284 | 0.0082 | -0.1526 | 1455.421 | 9.179689 | 57 |
| 96 | m. 5 | Phospholipid (PP) | 0.00240 | -0.0021 | 0.0564 | 377.2674 | 11.743289 | 691 |
| 97 | m. 5 | Lipid (Li) | 0.00167 | 0.0404 | -0.0329 | 279.2328 | 11.961267 | 834 |
| 102 | m. 5 | Lipid (Li) | 0.00128 | 0.0511 | -0.0726 | 281.2485 | 13.161200 | 884 |
| 124 | m. 5 | Peptide (Pe) | 0.00173 | 0.0786 | -0.1331 | 1099.6413 | 16.987733 | 583 |
| 49 | m. 6 | Alkaloid (Al) | 0.00323 | 0.0532 | 0.1335 | 445.369 | 5.779267 | 829 |
| 98 | m. 6 | Saponin (Sa) | 0.00015 | -0.0269 | -0.0464 | 607.2933 | 12.283867 | 483 |
| 107 | m. 6 | Phospholipid (PP) | -0.00005 | -0.0122 | -0.0705 | 593.2777 | 14.143022 | 468 |
| 108 | m. 6 | Sterol (St) | -0.00004 | -0.0533 | -0.0551 | 621.3084 | 14.628844 | 496 |
| 110 | m. 6 | Peptide (Pe) | 0.00014 | -0.0388 | -0.0504 | 635.3816 | 15.485111 | 601 |
| 121 | m. 6 | Phospholipid (PP) | -0.00057 | -0.0485 | -0.0593 | 607.2933 | 16.585889 | 483 |
| 126 | m. 6 | Phospholipid (PP) | -0.00031 | -0.0179 | -0.0777 | 621.3084 | 17.422600 | 496 |
| 132 | m. 6 | Peptide (Pe) | -0.00042 | -0.0410 | -0.1018 | 635.3816 | 18.238778 | 601 |
| 66 | m. 7 | Saponin (Sa) | -0.00238 | 0.0110 | -0.0479 | 1087.5707 | 6.218867 | 525 |
| 68 | m. 7 | Saponin (Sa) | 0.00025 | -0.0381 | 0.0862 | 943.529 | 6.721333 | 561 |
| 74 | m. 7 | Saponin (Sa) | 0.00259 | -0.0129 | 0.0691 | 585.4277 | 7.205911 | 731 |
| 75 | m. 7 | Saponin (Sa) | 0.00227 | -0.0067 | 0.0576 | 907.5367 | 7.203356 | 591 |
| 76 | m. 7 | Saponin (Sa) | 0.00190 | 0.0480 | 0.0452 | 1069.5625 | 7.218822 | 526 |
| 79 | m. 7 | Saponin (Sa) | -0.00375 | -0.0688 | -0.0314 | 923.5027 | 7.572822 | 544 |
| 81 | m. 7 | Phospholipid (PP) | 0.00063 | 0.0602 | -0.0630 | 277.2167 | 8.279133 | 782 |
| 82 | m. 7 | Phospholipid (PP) | 0.00058 | 0.0288 | -0.0403 | 317.209 | 8.321022 | 659 |
| 31 | m. 8 | Phen. glyco. (Pg) | 0.00117 | -0.0678 | -0.1842 | 684.2517 | 5.347200 | 368 |
| 40 | m. 8 | Saponin (Sa) | 0.00123 | 0.1758 | 0.0983 | 1119.5627 | 5.806133 | 503 |
| 43 | m. 8 | Saponin (Sa) | 0.00055 | 0.0050 | 0.0909 | 973.504 | 5.631911 | 518 |
| 46 | m. 8 | Saponin (Sa) | 0.00093 | 0.0569 | -0.2069 | 989.4993 | 5.818644 | 505 |
| 47 | m. 8 | Saponin (Sa) | 0.00173 | -0.1150 | 0.0430 | 843.4409 | 5.873844 | 523 |
| 52 | m. 8 | Saponin (Sa) | 0.00121 | -0.0305 | 0.1098 | 973.504 | 6.013467 | 518 |
| 56 | m. 8 | Saponin (Sa) | 0.00285 | -0.0381 | 0.0958 | 827.4455 | 6.058733 | 538 |
| 16 | m. 9 | Peptide (Pe) | -0.00141 | -0.0005 | 0.0386 | 141.0881 | 3.049089 | 624 |
| 19 | m. 9 | Alkaloid (Al) | 0.00186 | 0.0813 | 0.0131 | 422.1973 | 4.658444 | 467 |
| 72 | m. 9 | Peptide (Pe) | -0.00253 | 0.0377 | -0.0505 | 1170.1459 | 7.037956 | 125 |
| 73 | m. 9 | Saponin (Sa) | -0.00558 | -0.0807 | 0.0211 | 402.1992 | 7.065644 | 495 |
| 125 | m. 9 | Peptide (Pe) | -0.00057 | 0.1016 | 0.1027 | 584.4235 | 16.991733 | 725 |
| 181 | m. 9 | Halogen (Ha) | -0.00116 | 0.0197 | -0.0283 | 127.1233 | 27.473089 | 970 |
| 117 | m. 10 | Peptide (Pe) | 0.00509 | 0.0172 | 0.0481 | 595.2837 | 16.515511 | 477 |
| 118 | m. 10 | Peptide (Pe) | 0.00425 | 0.0036 | 0.0906 | 613.4849 | 16.515178 | 790 |
| 137 | m. 10 | Sterol (St) | 0.00234 | -0.0163 | 0.0588 | 335.2595 | 18.765067 | 774 |
| 138 | m. 10 | Phospholipid (PP) | 0.00259 | -0.0120 | 0.0542 | 613.4849 | 18.837933 | 790 |
| 150 | m. 10 | Phospholipid (PP) | 0.00178 | -0.0083 | 0.0982 | 775.535 | 20.840600 | 690 |
| 39 | m. 11 | Saponin (Sa) | 0.00214 | -0.1306 | 0.0287 | 527.2482 | 5.672422 | 471 |
| 176 | m. 11 | Phospholipid (PP) | 0.00557 | -0.0410 | 0.0270 | 861.4798 | 26.956867 | 557 |
| 178 | m. 11 | Phospholipid (PP) | 0.00338 | -0.0390 | -0.0627 | 835.5451 | 27.192933 | 652 |
| 179 | m. 11 | Phospholipid (PP) | 0.00517 | -0.0743 | -0.0850 | 834.5424 | 27.162333 | 650 |
| 20 | m. 12 | Saponin (Sa) | 0.00021 | 0.0792 | 0.1608 | 409.1823 | 4.755400 | 313 |
| 21 | m. 12 | Phen. glyco. (Pg) | -0.00217 | -0.0762 | -0.1660 | 481.2056 | 4.897222 | 427 |
| 24 | m. 12 | Peptide (Pe) | 0.00115 | 0.0097 | 0.1753 | 374.1461 | 5.014222 | 390 |
| 25 | m. 12 | Saponin (Sa) | 0.00127 | 0.0686 | -0.1764 | 475.2191 | 5.100889 | 461 |
| 48 | m. 12 | Peptide (Pe) | 0.00290 | 0.0146 | -0.1820 | 728.3151 | 5.887111 | 433 |
| 60 | m. 12 | Saponin (Sa) | 0.00040 | -0.1856 | 0.3557 | 959.5744 | 6.105156 | 599 |
| 116 | m. 12 | Peptide (Pe) | 0.00237 | -0.0299 | 0.0653 | 959.5744 | 16.460289 | 599 |
| 147 | m. 12 | Peptide (Pe) | 0.00043 | 0.0889 | 0.0042 | 965.6208 | 20.478333 | 643 |
| 26 | m. 13 | Phen. glyco. (Pg) | -0.00424 | 0.1468 | 0.0130 | 683.1501 | 5.126178 | 215 |
| 51 | m. 13 | Saponin (Sa) | 0.00026 | -0.0548 | 0.0378 | 915.4514 | 5.947444 | 659 |
| 61 | m. 13 | Saponin (Sa) | 0.00113 | 0.0149 | -0.0740 | 351.2153 | 6.255356 | 613 |
| 88 | m. 13 | Lipid (Li) | 0.00097 | 0.0176 | -0.0429 | 383.2564 | 9.921756 | 669 |
| 105 | m. 13 | Lipid (Li) | -0.00173 | -0.0380 | -0.0770 | 613.4839 | 13.432956 | 790 |
| 158 | m. 13 | Phospholipid (PP) | 0.00100 | 0.0318 | -0.0148 | 816.6582 | 21.926822 | 806 |
| 145 | m. 14 | Peptide (Pe) | 0.00156 | 0.0153 | 0.2906 | 939.5958 | 19.884289 | 634 |
| 152 | m. 14 | Phospholipid (PP) | -0.00445 | 0.1972 | -0.1885 | 941.6201 | 21.136822 | 659 |
| 156 | m. 14 | Sterol (St) | -0.00212 | -0.0755 | 0.1265 | 427.3947 | 21.699578 | 924 |

**TABLE A2.** Compounds and coefficients from lasso regression allowing for possible effects of all individual compounds (as in Figure 2) as well as all pairwise interactions among compounds. Listed here are interactions selected by lasso regression, using binomial and Gaussian regressions for survival and adult weight, respectively (with units on those scales, as in Table A1). Results for development time are not shown here as none of the potential interactions for development time were selected by lasso regression.

| Survival regression |  | Weight regression |  |  |  |
| --- | --- | --- | --- | --- | --- |
| Compounds | Coefficient | Compounds | Coefficient | Compounds | Coefficient |
| cpd.9 | -0.080 | cpd.25 | -0.085 | cpd.43 x cpd.47 | -0.030 |
| cpd.122 | 0.067 | cpd.49 | 0.0081 | cpd.49 x cpd.138 | 0.051 |
| cpd.154 | 0.0078 | cpd.5 x cpd.7 | -0.10 | cpd.55 x cpd.79 | -0.025 |
| cpd.5 x cpd.6 | -0.042 | cpd.5 x cpd.9 | -0.059 | cpd.60 x cpd.65 | 0.016 |
| cpd.5 x cpd.7 | -0.0095 | cpd.8 x cpd.11 | -0.018 | cpd.60 x cpd.79 | -0.048 |
| cpd.6 x cpd.34 | -0.024 | cpd.12 x cpd.152 | 0.019 | cpd.60 x cpd.132 | -0.080 |
| cpd.6 x cpd.79 | -0.11 | cpd.14 x cpd.91 | 0.11 | cpd.65 x cpd.133 | -0.22 |
| cpd.7 x cpd.133 | 0.078 | cpd.18 x cpd.73 | -0.18 | cpd.85 x cpd.161 | 0.16 |
| cpd.19 x cpd.54 | -0.0063 | cpd.18 x cpd.79 | 0.0048 | cpd.86 x cpd.141 | 0.052 |
| cpd.21 x cpd.60 | -0.038 | cpd.19 x cpd.85 | -0.024 | cpd.88 x cpd.118 | 0.063 |
| cpd.45 x cpd.54 | 0.00024 | cpd.19 x cpd.124 | 0.013 | cpd.91 x cpd.122 | -0.062 |
| cpd.48 x cpd.142 | 0.019 | cpd.23 x cpd.24 | -0.011 | cpd.95 x cpd.175 | -0.011 |
| cpd.61 x cpd.82 | -0.034 | cpd.25 x cpd.60 | 0.086 | cpd.105 x cpd.125 | 0.024 |
| cpd.62 x cpd.79 | -0.010 | cpd.25 x cpd.117 | -0.019 | cpd.105 x cpd.152 | -0.090 |
| cpd.83 x cpd.164 | -0.017 | cpd.25 x cpd.145 | -0.078 | cpd.117 x cpd.149 | -0.10 |
| cpd.99 x cpd.109 | -0.039 | cpd.32 x cpd.66 | -0.37 | cpd.117 x cpd.156 | 0.047 |
| cpd.152 x cpd.174 | -0.18 | cpd.34 x cpd.94 | 0.00025 | cpd.121 x cpd.140 | 0.25 |
|  |  | cpd.35 x cpd.49 | -0.062 | cpd.125 x cpd.148 | 0.0035 |
|  |  | cpd.41 x cpd.179 | 0.144 | cpd.124 x cpd.176 | 0.13 |

**FIGURE A1.**
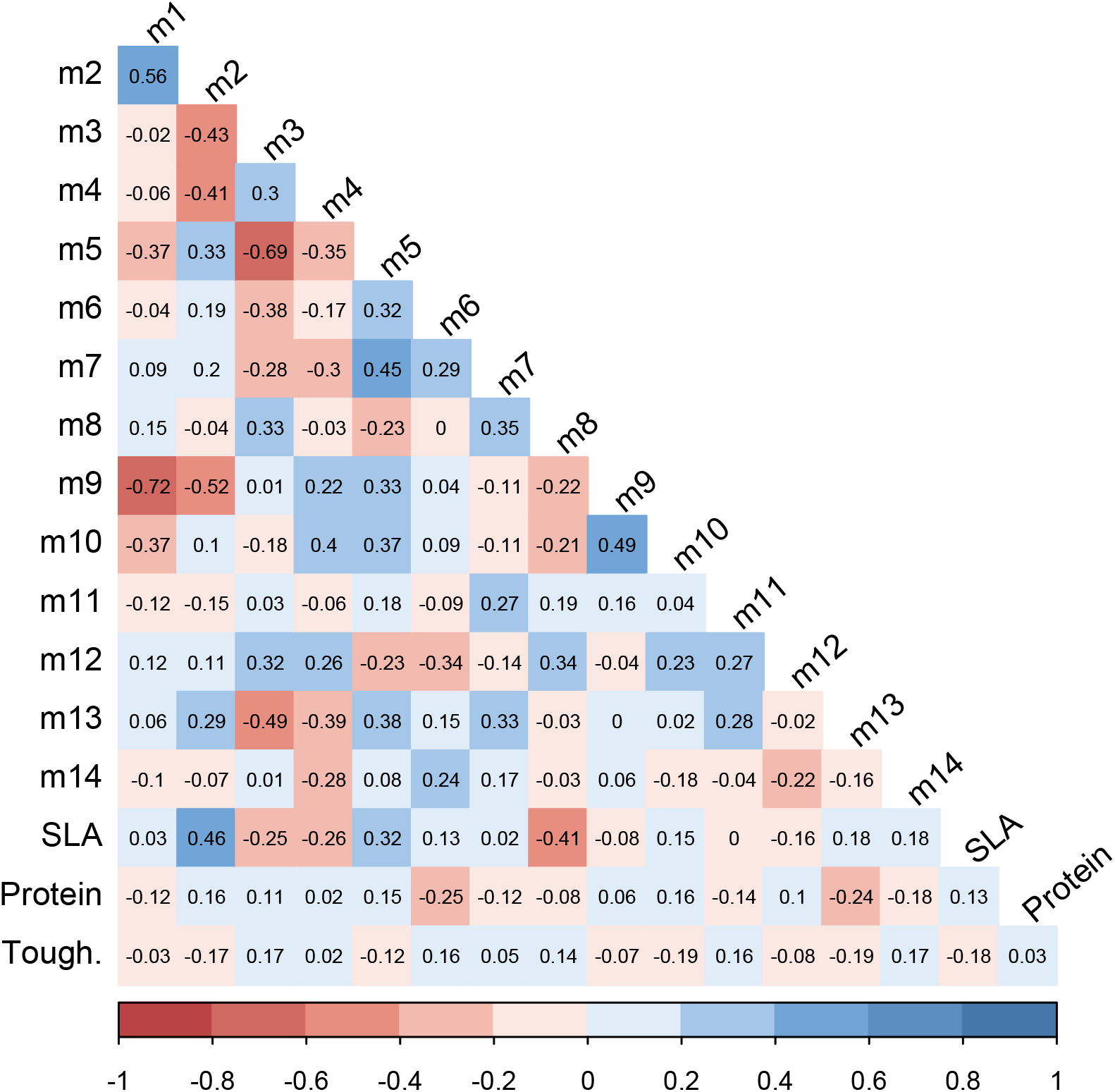
Pairwise correlations between phytochemical modules (represented by eigenvectors m1, m2, etc.) and plant traits specific leaf area (SLA), protein and leaf toughness. Values shown are Pearson product-moment correlation coefficients.

**FIGURE A2a.**
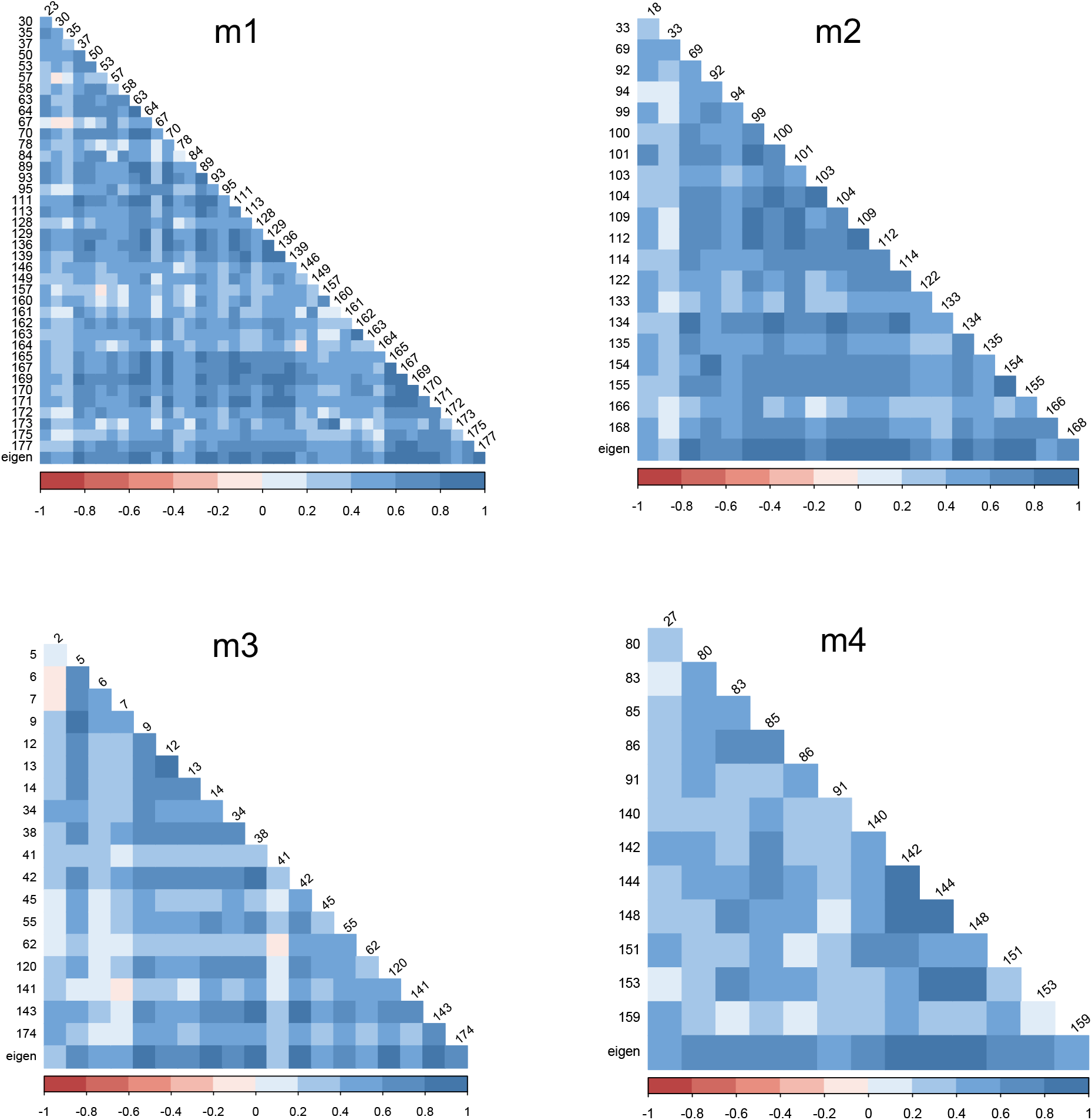
Pairwise correlations (as in Figure A1), but among compounds within modules m1 to m4. The bottom row in each graph shows correlations among individual compounds and the eigenvector used in other analyses for the given module.

**FIGURE A2b.**
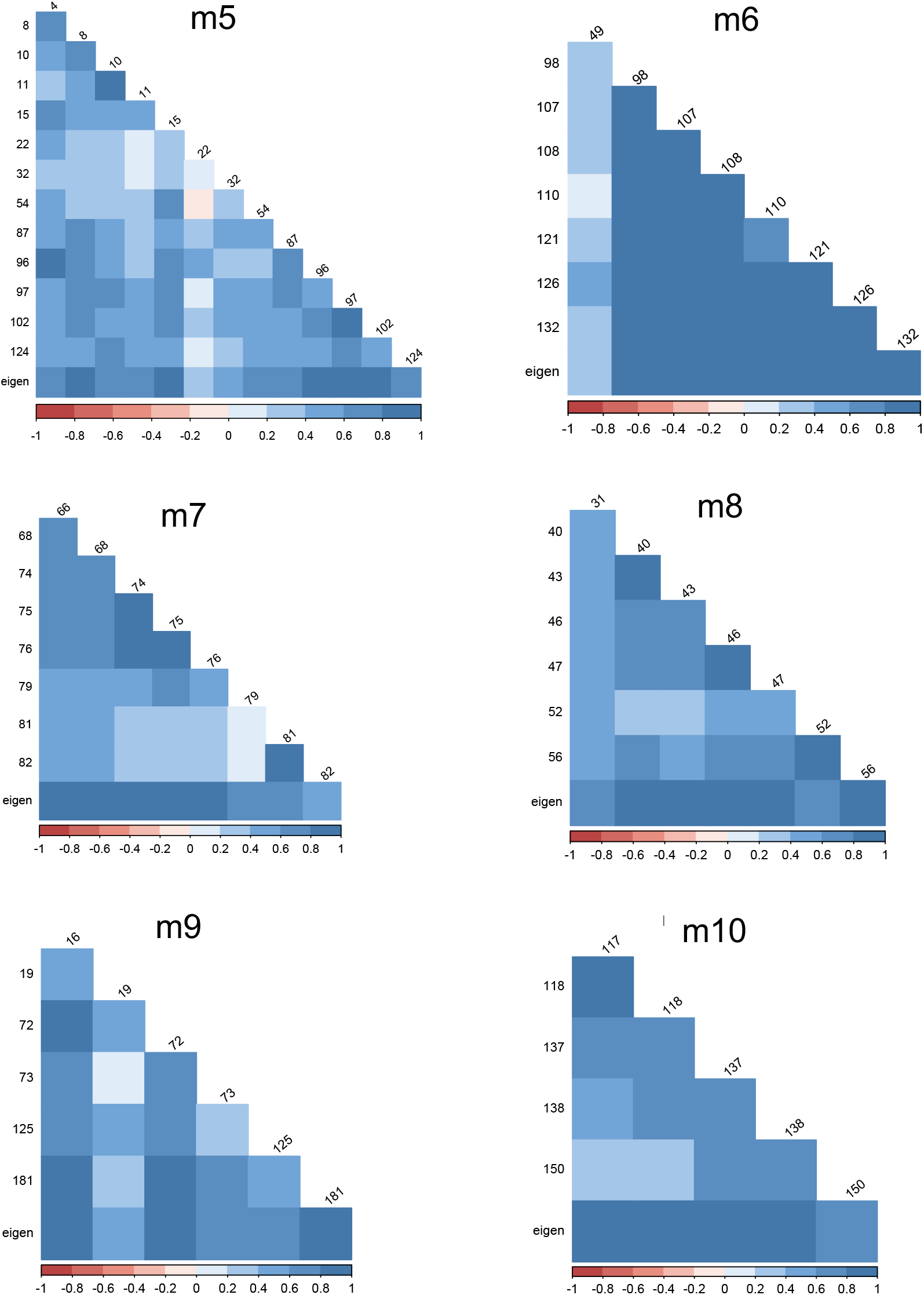
Pairwise correlations (as in Figure A1), but among compounds within modules m5 to m10. The bottom row in each graph shows correlations among individual compounds and the eigenvector used in other analyses for the given module.

**FIGURE A2c.**
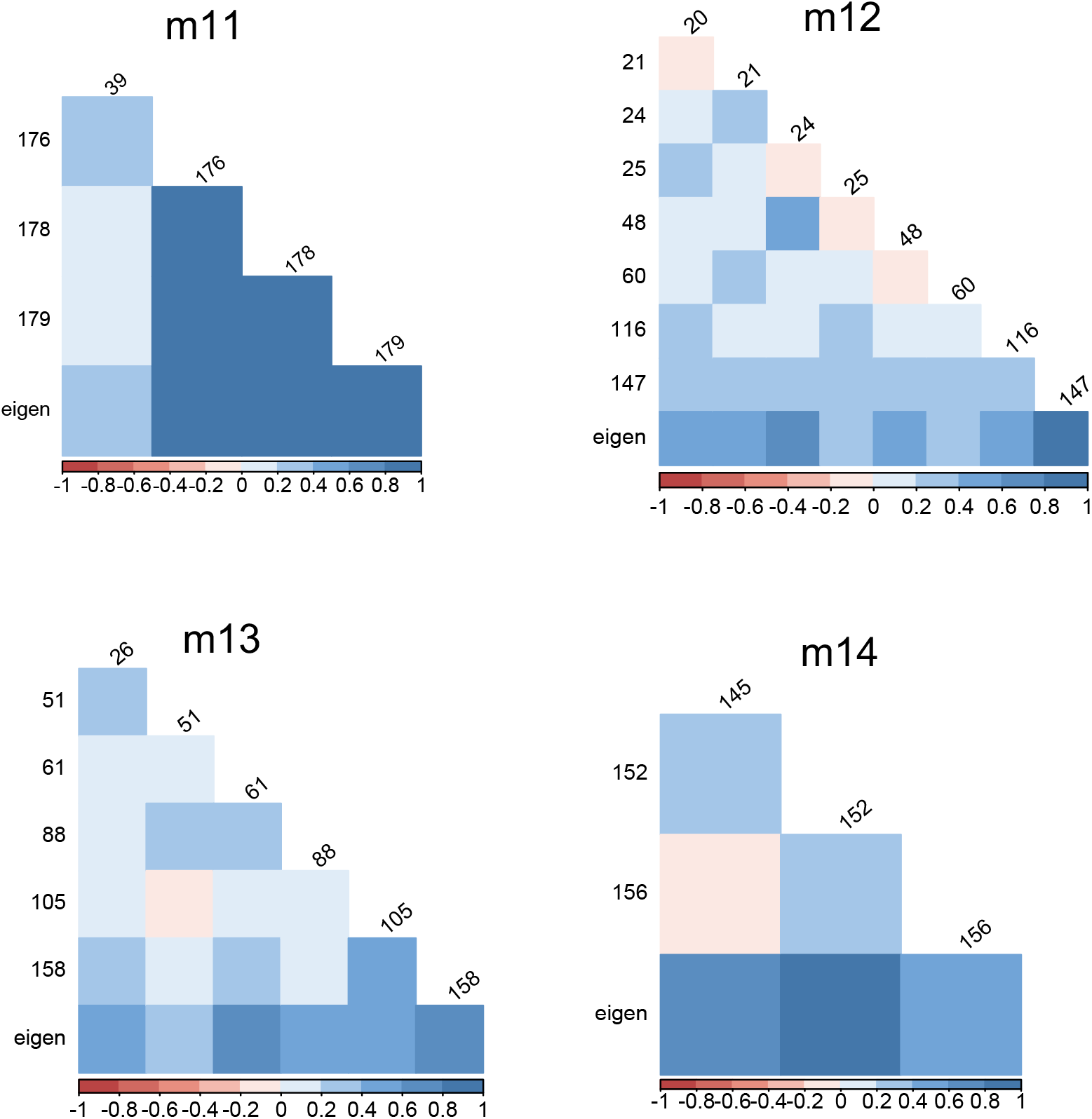
Pairwise correlations (as in Figure A1), but among compounds within modules m11 to m14. The bottom row in each graph shows correlations among individual compounds and the eigenvector used in other analyses for the given module.

**FIGURE A3.**
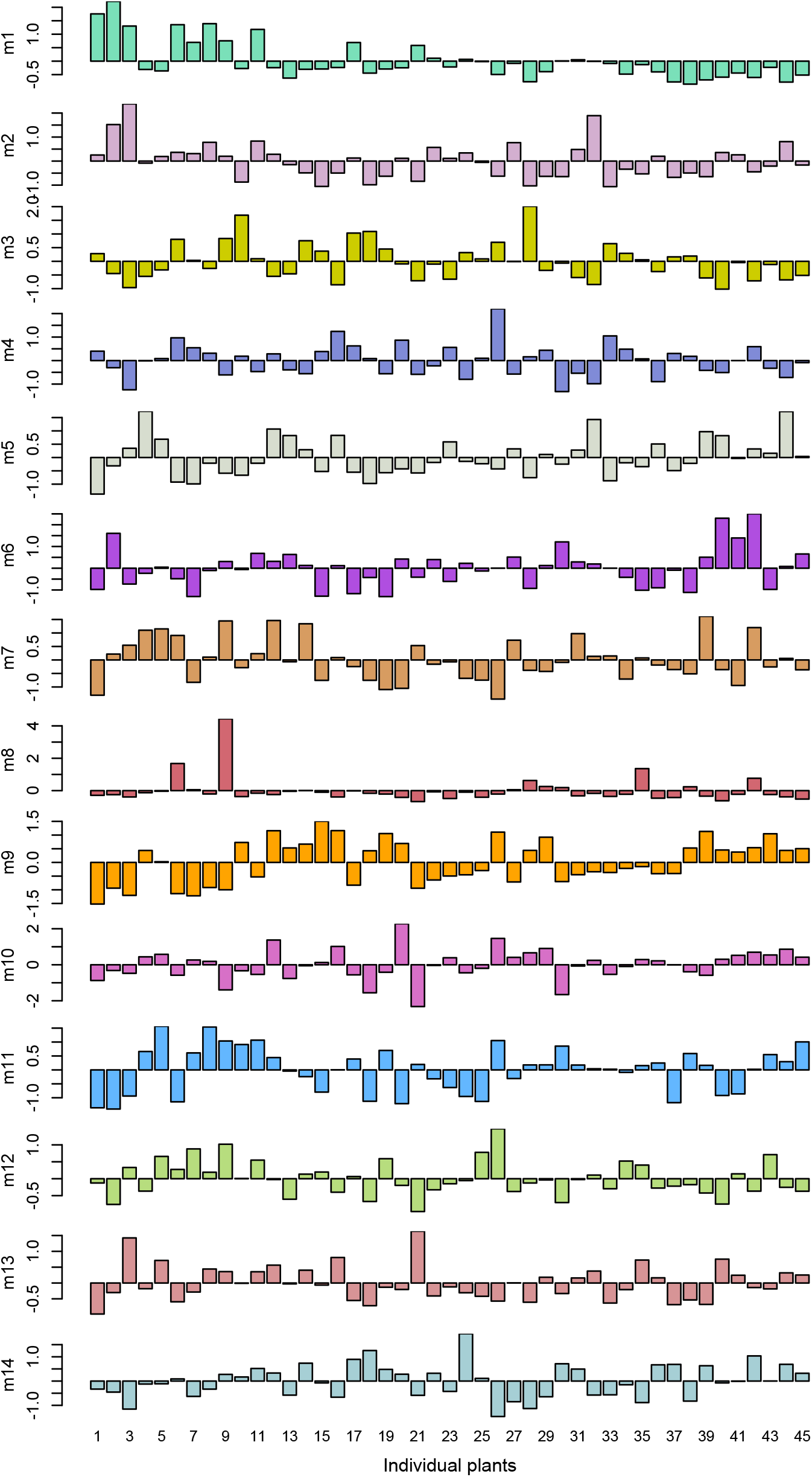
Visualization of variation among plants (bars) in phytochemical modules. Each bar is the average of z-scores for compounds comprising a given module. Colors correspond to modules as in Figure 1. The order of plants (along the x-axis) is arbitrary.

**FIGURE A4.**
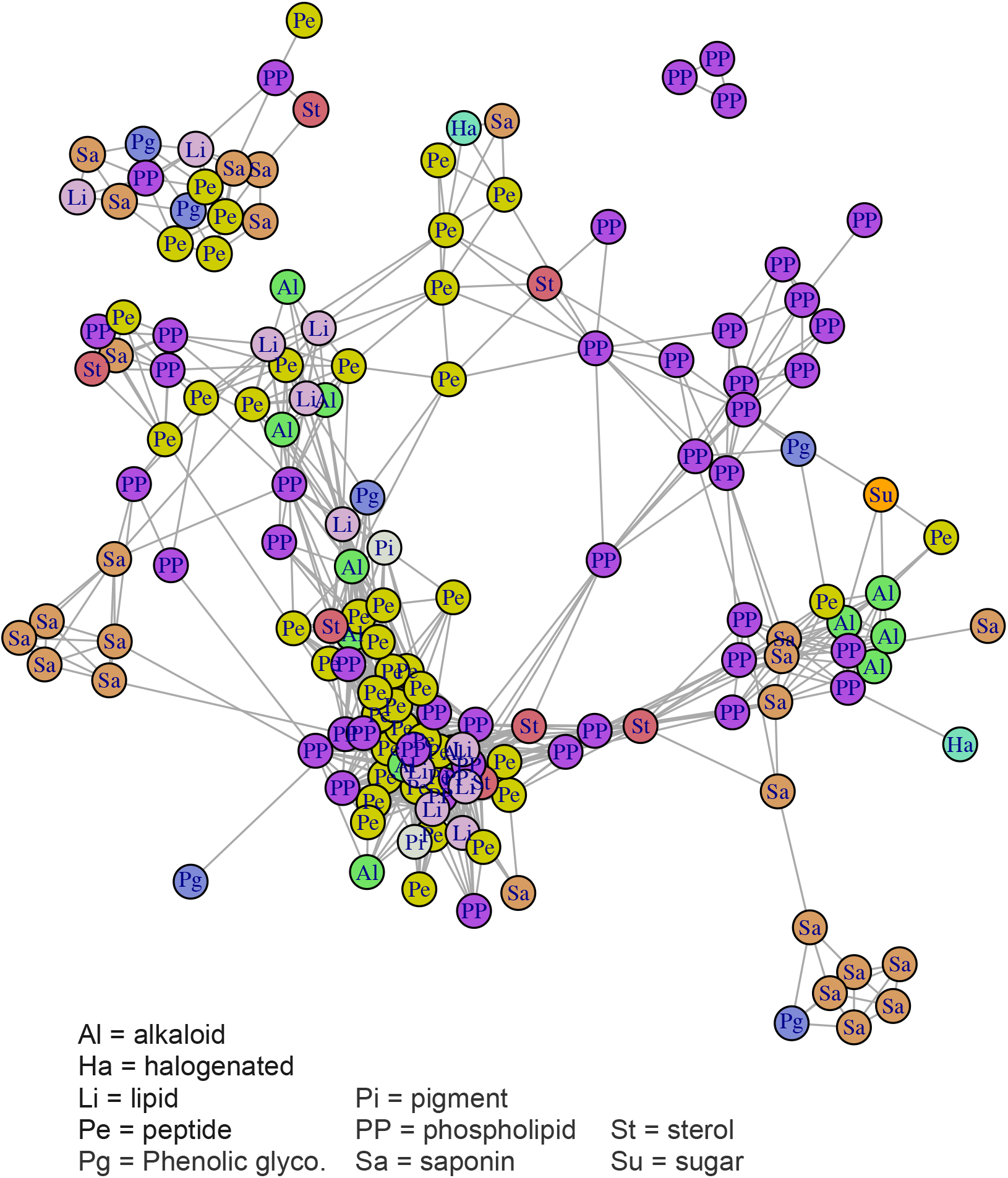
Illustration of correlational structure among compounds, as in Figure 2, but here color coded by compound class identities instead of by module assignment.

**FIGURE A5.**
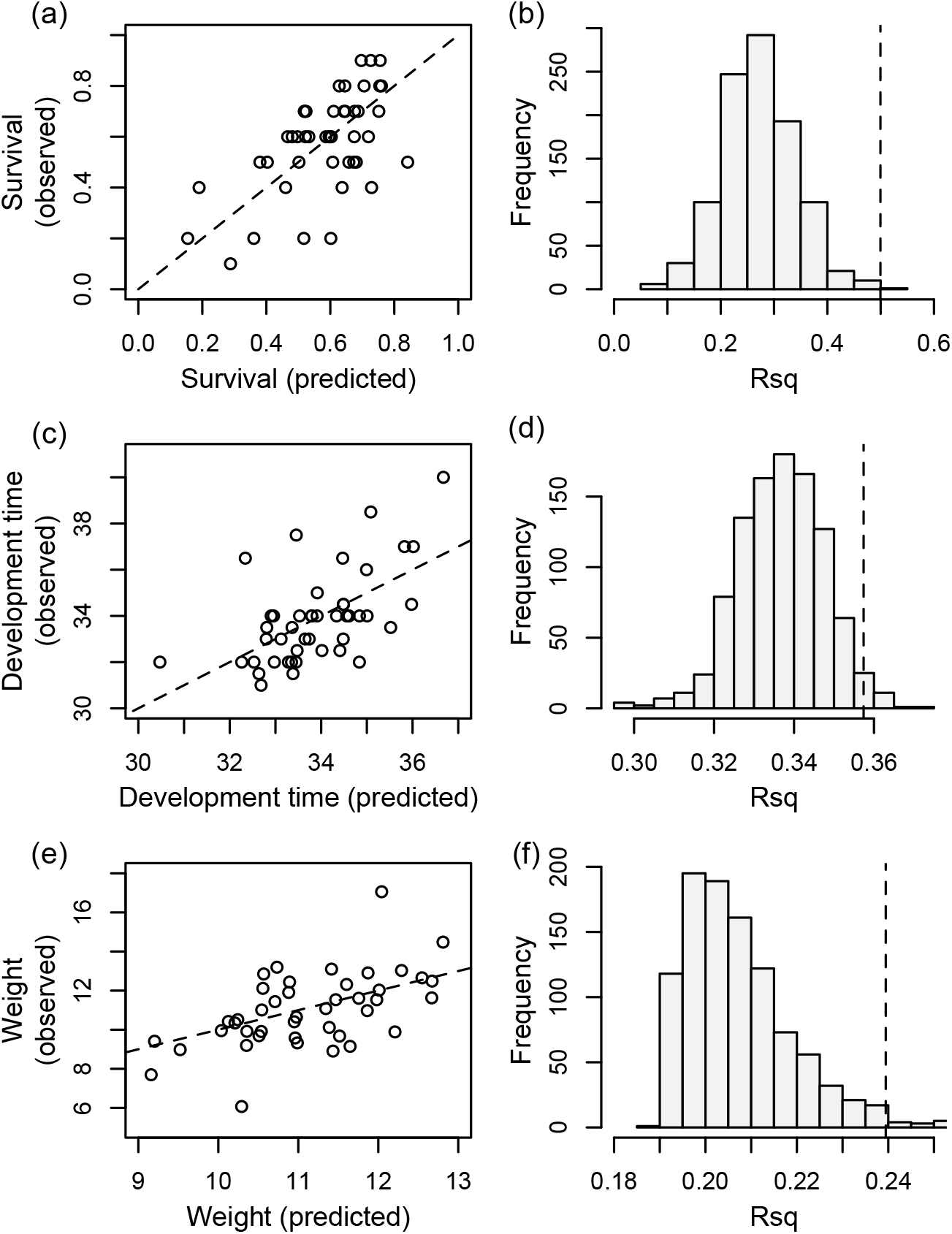
Results from cross-validation (a, c, e) and resampling analysis (b, d, f) of Bayesian regressions reported in Table 1. In the observed vs predicted plots (a, c, e), each dot represents the observed and predicted performance of caterpillars associated with individual plants, which was the level at which cross-validation was conducted (i.e., one plant was left out of each iteration). In the resampling analyses (b, d, f), “modules” were constructed based on randomly assembled collections of compounds selected to match the structure of the initial analyses. For example, the survival regressions included modules 2, 3, 9, 10, and 11 (Table 1). Because m2 includes 21 compounds, for one iteration of the resampling analysis 21 compounds were randomly selected, from which the first eigenvector was taken for use in the analysis (and same for the other modules). From each of 1000 randomly-created sets of modules, R^2^ values were retained and summarized in plots (b, d, f), along with dashed lines representing the R^2^ values of empirical collections of compounds and associated eigenvectors. In all cases, the empirical models outperformed all but a tiny fraction of simulated models. The fraction of the simulated models with R^2^ values greater than the empirical models are as follows: 0.007 (b), 0.027 (d), and 0.023 (f).

**FIGURE A6.**
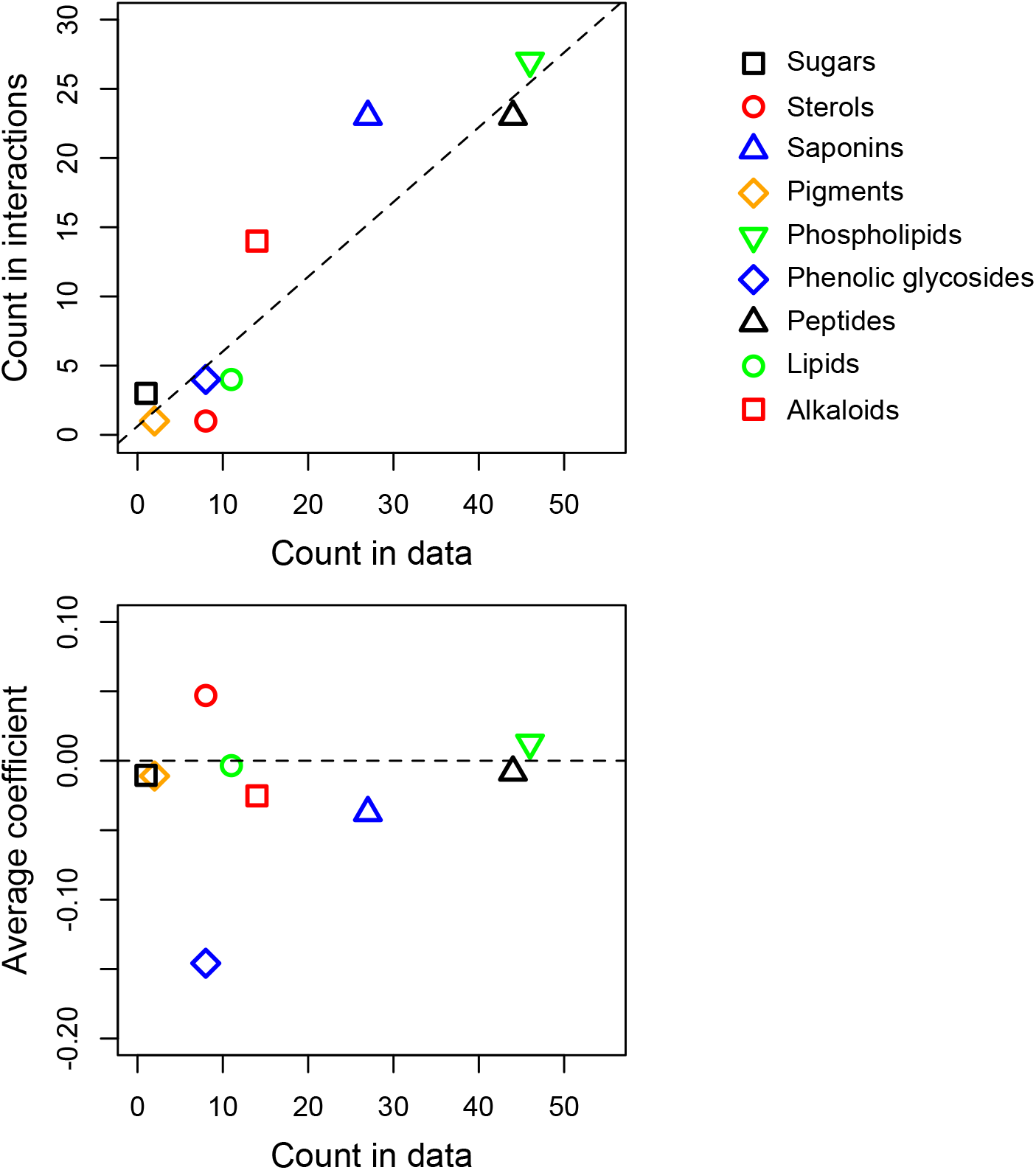
Additional details on interactions by compound class. Top panel shows the number of times that compounds of a certain class appear in pairwise interactions (as in appendix Table A2) as a function of the total numbers of those compounds in different classes (“Count in data”). Classes that are represented by a greater number of distinct compounds in the total dataset are, in general, more likely to be observed in pairwise interactions: the dotted line shows the expected frequency of observation given random sampling from the total pool of compounds. Saponins and alkaloids are overrepresented in interactions, appearing in 44 and 41% (respectively) more pairwise interactions than expected by chance. Bottom panel shows average interaction coefficients for all interactions involving compounds of particular classes: most pairwise interactions are relatively small in magnitude, with the exception of interactions involving phenolic glycosides, that tend to be strongly negative. For both panels, interactions are considered from analyses of caterpillar survival and weight; no interactions were detected for development time (Supporting Information Table A2).

**FIGURE A7.**
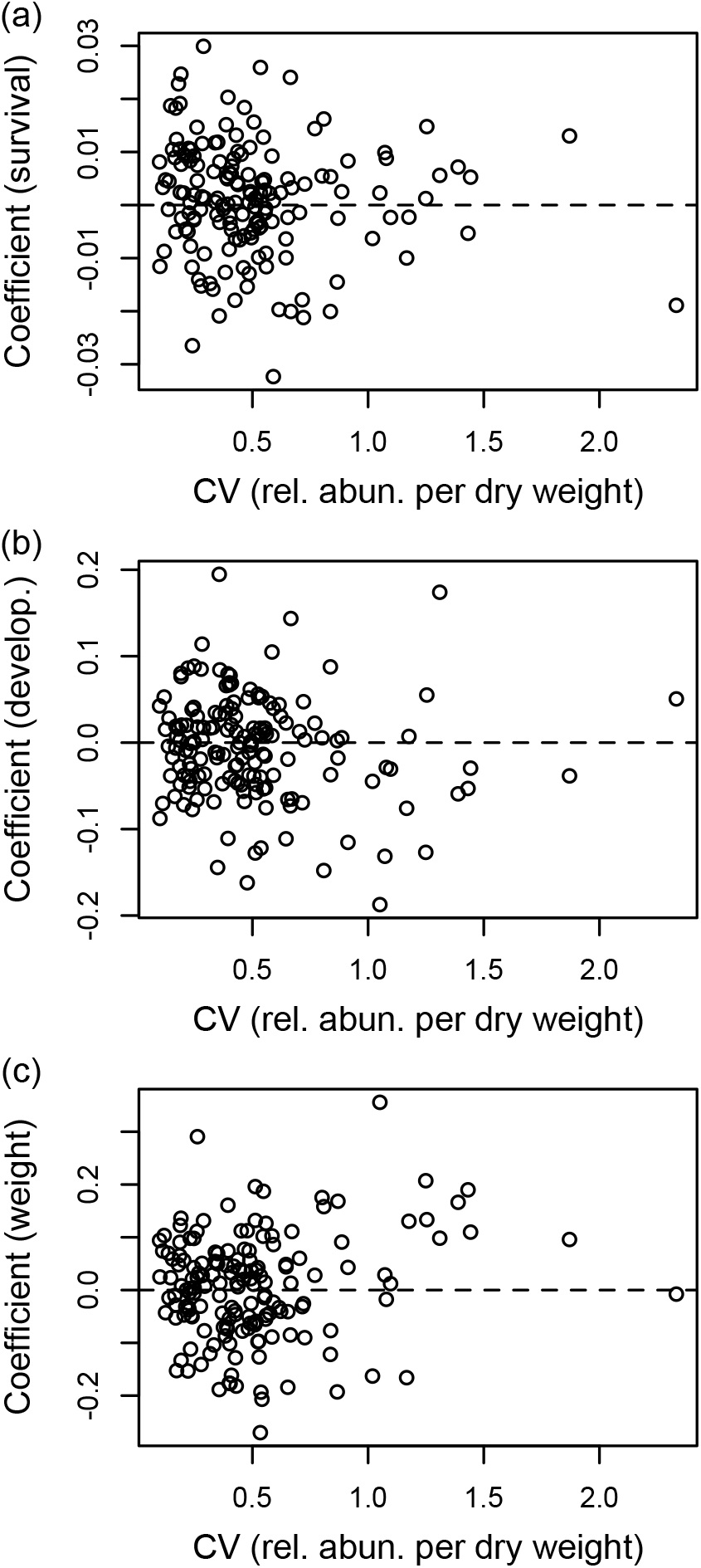
Compound specific effects vs coefficients of variation for individual compounds. The compound specific effects based on ridge regression are the same as those listed in Table A1.

**Figure A8.**
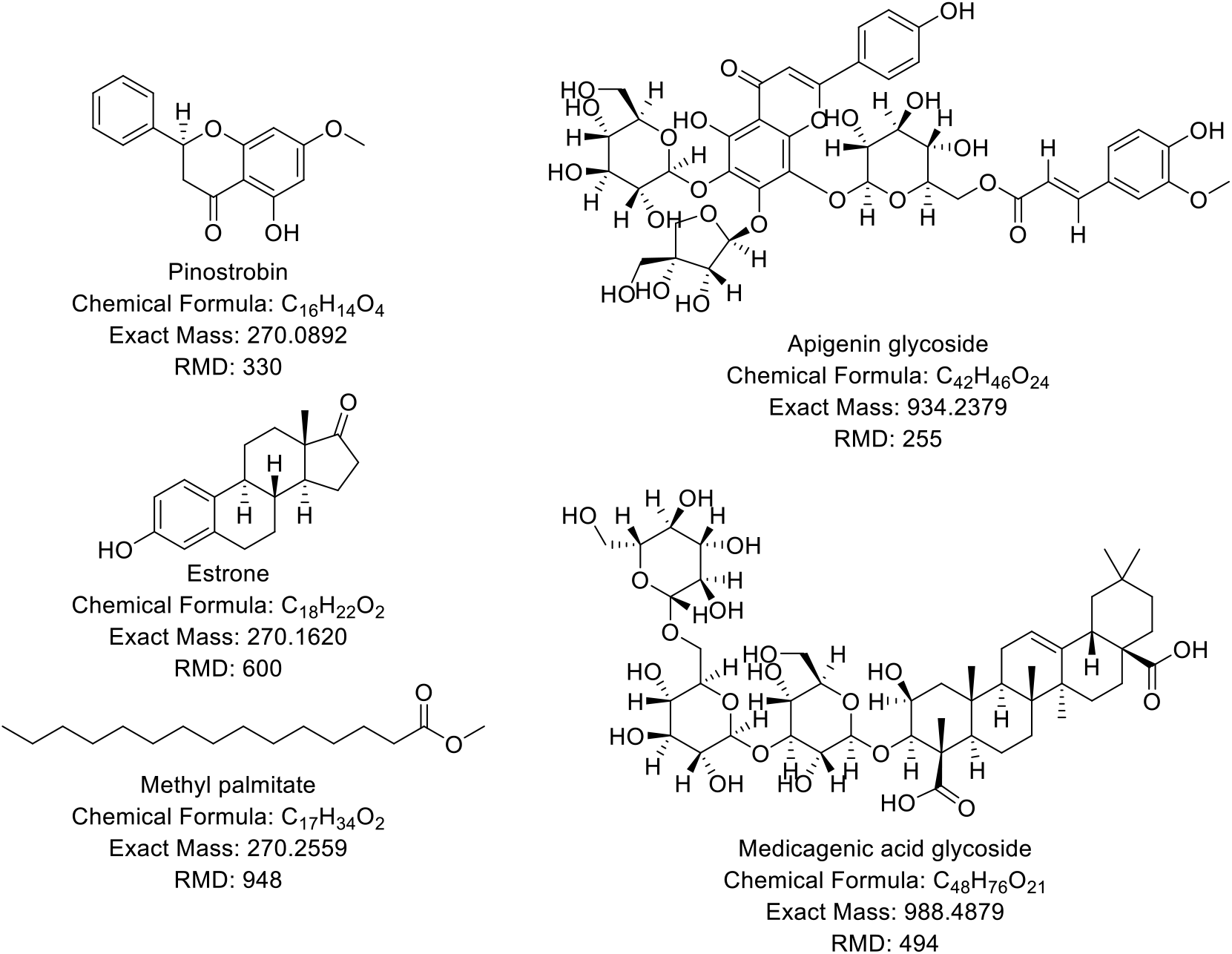
Relative mass defect of natural products. **Left:** A series of isobaric molecules (by nominal mass) which have increasing RMD as %H increases. **Right:** Characteristic metabolites from *M. sativa* which can be classified by RMD.

## REFERENCES

Agrawal, A. A., Petschenka, G., Bingham, R. A., Weber, M. G., & Rasmann, S. (2012). Toxic cardenolides: chemical ecology and coevolution of specialized plant--herbivore interactions. New Phytologist, 194(1), 28–45.

Balsevich, J. J., Bishop, G. G., & Deibert, L. K. (2009). Use of digitoxin and digoxin as internal standards in HPLC analysis of triterpene saponin-containing extracts. Phytochemical Analysis, 20(1), 38–49.

Behmer, S. T. (2009). Insect herbivore nutrient regulation. Annual Review of Entomology, 54, 165–187.

Berenbaum, M. (1983). Coumarins and caterpillars: a case for coevolution. Evolution, 37(1), 163–179.

Berenbaum, M. R. (1995). Turnabout is fair play - secondary roles for primary compounds. Journal of Chemical Ecology, 21, 925–940.

Brooks, S. P., & Gelman, A. (1998). General methods for monitoring convergence of iterative simulations. Journal of Computational and Graphical Statistics, 7(4), 434–455.

Carmona, D., Lajeunesse, M. J., & Johnson, M. T. J. (2011). Plant traits that predict resistance to herbivores. Functional Ecology, 25(2), 358–367.

Chaieb, I. (2010). Saponins as insecticides: a review. Tunisian Journal of Plant Protection, 5(1), 39–50.

Chaturvedi, S., Lucas, L. K., Nice, C. C., Fordyce, J. A., Forister, M. L., & Gompert, Z. (2018). The predictability of genomic changes underlying a recent host shift in Melissa blue butterflies. Molecular Ecology.

Delaney, N. J., & Chatterjee, S. (1986). Use of the bootstrap and cross-validation in ridge regression. Journal of Business & Economic Statistics, 4(2), 255–262.

Dyer, L A. (1995). Tasty generalists and nasty specialists - antipredator mechanisms in tropical lepidopteran larvae. Ecology, 76, 1483–1496.

Dyer, Lee A., Philbin, C. S., Ochsenrider, K. M., Richards, L. A., Massad, T. J., Smilanich, A. M., … Jeffrey, C. S. (2018). Modern approaches to study plant–insect interactions in chemical ecology. Nature Reviews Chemistry, 1. https://doi.org/10.1038/s41570-018-0009-7

Einstein, A. (1905). Ist die trägheit eines körpers von seinem energieinhalt abhängig? Annalen Der Physik, 323(13), 639–641.

Ekanayaka, E. A. P., Celiz, M. D., & Jones, A. D. (2015). Relative mass defect filtering of mass spectra: a path to discovery of plant specialized metabolites. Plant Physiology, 167, 1221– 1232. https://doi.org/10.1104/pp.114.251165

Erbilgin, N. (2018). Phytochemicals as mediators for host range expansion of a native invasive forest insect herbivore. New Phytologist.

Feeny, P., Rosenthal, G. A., & Berenbaum, M. R. (1992). The evolution of chemical ecology: contributions from the study of herbivorous insects. Herbivores: Their Interactions with Secondary Plant Metabolites, 2, 1–44.

Fordyce, J. A., & Nice, C. C. (2008). Antagonistic, stage-specific selection on defensive chemical sequestration in a toxic butterfly. Evolution, 62, 1610–1617. https://doi.org/10.1111/j.1558-5646.2008.00388.x

Forister, M. L., Nice, C. C., Fordyce, J. A., & Gompert, Z. (2009). Host range evolution is not driven by the optimization of larval performance: the case of *Lycaeides melissa* (Lepidoptera: Lycaenidae) and the colonization of alfalfa. Oecologia, 160, 551–561. https://doi.org/10.1007/s00442-009-1310-4

Forister, M. L., Scholl, C. F., Jahner, J. P., Wilson, J. S., Fordyce, J. A., Gompert, Z., … Nice, C. C. (2012). Specificity, rank preference and the colonization of a non-native host plant by the Melissa blue butterfly. Oecologia, DOI: 10.10.

Friedman, J., Hastie, T., Simon, N., & Tibshirani, R. (2016). Lasso and elastic-net regularized generalized linear models. R-package version 2.0-5. 2016.

Gelman, A., Rubin, D. B., & others. (1992). Inference from iterative simulation using multiple sequences. Statistical Science, 7(4), 457–472.

Glassmire, A. E., Philbin, C., Richards, L. A., Jeffrey, C. S., Snook, J. S., & Dyer, L. A. (2019). Proximity to canopy mediates changes in the defensive chemistry and herbivore loads of an understory tropical shrub, *Piper kelleyi*. Ecology Letters, 22(2), 332–341.

Gompert, Z., Brady, M., Chalyavi, F., Saley, T. C., Philbin, C. S., Tucker, M. J., … Lucas, L. K. (2019). Genomic evidence of genetic variation with pleiotropic effects on caterpillar fitness and plant traits in a model legume. Molecular Ecology, 28(12), 2967–2985.

Gompert, Z., Jahner, J. P., Scholl, C. F., Wilson, J. S., Lucas, L. K., Soria-Carrasco, V., … Forister, M. L. (2015). The evolution of novel host use is unlikely to be constrained by trade-offs or a lack of genetic variation. Molecular Ecology, 24, 2777–2793.

Harrison, J. G., Gompert, Z., Fordyce, J. A., Buerkle, C. A., Grinstead, R., Jahner, J. P., … Forister, M. L. (2016). The many dimensions of diet breadth: phytochemical, genetic, behavioral, and physiological perspectives on the interaction between a native herbivore and an exotic host. PloS One, 11, e0147971.

Harrison, J. G., Philbin, C. S., Gompert, Z., Forister, G. W., Hernandez-Espinoza, L., Sullivan, B. W., … others. (2018). Deconstruction of a plant-arthropod community reveals influential plant traits with nonlinear effects on arthropod assemblages. Functional Ecology, 32(5), 1317–1328.

Hättenschwiler, S., & Vitousek, P. M. (2000). The role of polyphenols in terrestrial ecosystem nutrient cycling. Trends in Ecology & Evolution, 15(6), 238–243.

Hunter, M. D. (2016). *The phytochemical landscape: linking trophic interactions and nutrient dynamics*. Princeton University Press.

Jansen, J. J., Allwood, J. W., Marsden-Edwards, E., van der Putten, W. H., Goodacre, R., & van Dam, N. M. (2009). Metabolomic analysis of the interaction between plants and herbivores. Metabolomics, 5(1), 150.

Jorge, T. F., Mata, A. T., & António, C. (2016). Mass spectrometry as a quantitative tool in plant metabolomics. Phil. Trans. R. Soc. A, 374(2079), 20150370.

Langfelder, P., & Horvath, S. (2008). WGCNA: an R package for weighted correlation network analysis. BMC Bioinformatics, 9(1), 559.

Levin, D. A. (1976). The chemical defenses of plants to pathogens and herbivores. Annual Review of Ecology and Systematics, 7(1), 121–159.

Maag, D., Erb, M., & Glauser, G. (2015). Metabolomics in plant-herbivore interactions: challenges and applications. Entomologia Experimentalis et Applicata, 157(1), 18–29.

Macel, M., van Dam, N. M., & Keurentjes, J. J. B. (2010). Metabolomics: the chemistry between ecology and genetics. Molecular Ecology Resources, 10(4), 583–593.

Mattila, H. R., & Otis, G. W. (2003). A comparison of the host preference of monarch butterflies (*Danaus plexippus*) for milkweed (*Asclepias syriaca*) over dog-strangler vine (*Vincetoxicum rossicum*). Entomologia Experimentalis et Applicata, 107, 193–199.

Ogutu, J. O., Schulz-Streeck, T., & Piepho, H.-P. (2012). Genomic selection using regularized linear regression models: ridge regression, lasso, elastic net and their extensions. In BMC proceedings (Vol. 6, p. S10).

Philbin, C. S., & Forister, M. L. (n.d.). Clustering and classification of phytochemicals from an experimental system using Bayesian model clustering.

Plummer, M., & others. (2003). JAGS: A program for analysis of Bayesian graphical models using Gibbs sampling. In Proceedings of the 3rd international workshop on distributed statistical computing (Vol. 124).

Prince, E. K., & Pohnert, G. (2010). Searching for signals in the noise: metabolomics in chemical ecology. Analytical and Bioanalytical Chemistry, 396(1), 193–197.

RCoreDevelopmentTeam. (2016). R: A Language and Environment for Statistical Computing. R Foundation for Statistical Computing. https://doi.org/10.1007/978-3-540-74686-7

Richards, L. A., Dyer, L. A., Smilanich, A. M., & Dodson, C. D. (2010). Synergistic effects of amides from two *Piper* species on generalist and specialist herbivores. Journal of Chemical Ecology, 36(10), 1105–1113.

Salazar, D., Lokvam, J., Mesones, I., Pilco, M. V., Zuñiga, J. M. A., de Valpine, P., & Fine, P. V. A. (2018). Origin and maintenance of chemical diversity in a species-rich tropical tree lineage. Nature Ecology & Evolution, 2(6), 983–990.

Sardans, J., Penuelas, J., & Rivas-Ubach, A. (2011). Ecological metabolomics: overview of current developments and future challenges. Chemoecology, 21(4), 191–225.

Seigler, D., & Price, P. W. (1976). Secondary compounds in plants: primary functions. The American Naturalist, 110(971), 101–105.

Smilanich, A. M., Fincher, R. M., & Dyer, L. A. (2016). Does plant apparency matter? Thirty years of data provide limited support but reveal clear patterns of the effects of plant chemistry on herbivores. New Phytologist, 210(3), 1044–1057.

Smith, C. A., O’Maille, G., Want, E. J., Qin, C., Trauger, S. A., Brandon, T. R., … Siuzdak, G. (2005). METLIN: a metabolite mass spectral database. Ther Drug Monit, 27, 747–751. Retrieved from http://www.ncbi.nlm.nih.gov/pubmed/16404815

Wu, S., Wilson, A. E., Chang, L., & Tian, L. (2019). Exploring the phytochemical landscape of the early-diverging flowering plant *Amborella trichopoda* Baill. Molecules, 24(21), 3814.

Zalucki, M. P., Brower, L. P., & Alonso-M, A. (2001). Detrimental effects of latex and cardiac glycosides on survival and growth of first-instar monarch butterfly larvae *Danaus plexippus* feeding on the sandhill milkweed *Asclepias humistrata*. Ecological Entomology, 26(2), 212–224. https://doi.org/10.1046/j.1365-2311.2001.00313.x

